# Secretion is a second fate for p62 and is coupled to KEAP1/NRF2 signaling

**DOI:** 10.64898/2026.10.07.757498

**Authors:** Koki Saito, Tomohiro Takahashi, Masashi Arakawa, Eiji Morita

**Affiliations:** Department of Biochemistry and Molecular Biology, Faculty of Agriculture and Life Science, Hirosaki University, 3 Bunkyo-cho, Hirosaki, Aomori 036-8561, Japan; Division of Biomolecular Function, Bioresources Science, United Graduate School of Agricultural Sciences, Iwate University, Morioka 020-0066, Japan

## Abstract

Cells degrade the autophagy receptor p62/sequestosome 1 (SQSTM1) in lysosomes. By knocking the HiBiT peptide into the endogenous SQSTM1 locus, we show that export is a second, quantitatively comparable fate. Specifically, p62 leaves cells continuously inside membrane-enclosed carriers at a rate similar to that of LC3B. Blocking lysosomal degradation amplifies release through an ATG5- and FIP200-dependent route, whereas a basal route persists in autophagy-deficient cells. Thus, p62 exits by at least two parallel pathways. Export requires PB1-mediated self-oligomerization but not LC3 binding, Kelch-like ECH-associated protein 1 (KEAP1) binding, or an intact ubiquitin-associated domain. Hence, oligomerization is the sorting determinant. KEAP1 is also released, and dominant-negative RAB27A retains both proteins inside cells and lowers nuclear factor erythroid 2-related factor 2 transcriptional output. Secretion is therefore a second outlet for the p62–KEAP1 module, and its interruption co-varies with a dampened oxidative stress response.

## Introduction

Macroautophagy (hereafter referred to as autophagy) is a bulk degradation system in which cytoplasmic constituents are sequestered by a double-membrane phagophore, which is enclosed within an autophagosome and delivered to the lysosome for hydrolysis (Dikic and Elazar, 2018; Mizushima and Komatsu, 2011). Beyond this degradative output, an alternative fate of the autophagic cargo has become increasingly apparent. In particular, under conditions in which lysosomal function is limited, autophagic membranes and their contents are transported to the plasma membrane and released into the extracellular space. This process is collectively termed secretory autophagy (Ponpuak et al., 2015). Leaderless cytosolic proteins, including interleukin-1β, are exported through this route (Dupont et al., 2011; Kimura et al., 2017), and the microtubule-associated protein 1 light chain 3 (LC3)-conjugation machinery directs the loading of selected cargo, particularly RNA-binding proteins, into extracellular vesicles (EV) in a process designated LC3-dependent EV loading and secretion (Leidal and Debnath, 2020; Leidal et al., 2020). Pharmacological or genetic inactivation of lysosomes triggers a coordinated secretory response that disposes of otherwise undegradable material, thereby preserving proteostasis (Solvik et al., 2022). Autophagy-derived EVs are biochemically distinct from canonical exosomes (Mao et al., 2025). Despite this progress, the identity of the cargo that traverses these routes, the relative contribution of each route, and the physiological consequences of the resulting extracellular pool remain largely unknown.

The p62/sequestosome 1 (SQSTM1) receptor is a prototypical selective autophagy receptor. Through its C-terminal ubiquitin-associated (UBA) domain, the receptor engages ubiquitylated substrates. Through its LC3-interacting region (LIR) motif, it engages LC3/gamma-aminobutyric acid type A receptor-associated proteins on the phagophore. In addition, p62/SQSTM1 self-oligomerizes through its N-terminal Phox and Bem1 (PB1) domain into helical filaments that nucleate micrometer-scale phase-separated p62 bodies (Itakura and Mizushima, 2011; Jakobi et al., 2020; Katsuragi et al., 2015; Sun et al., 2018). Accordingly, p62 is a cargo receptor and signaling hub, and its Kelch-like ECH-associated protein 1 (KEAP1)-interacting region (KIR) competes with nuclear factor erythroid 2-related factor 2 (NRF2) for the Kelch pocket of KEAP1. Phosphorylation of Ser349 within the KIR increases this affinity, thereby sequestering KEAP1, stabilizing NRF2, and inducing antioxidant and detoxifying gene programs (Ichimura et al., 2013; Komatsu et al., 2010; Yamamoto et al., 2018). As p62 is itself an autophagic substrate, its intracellular concentration reports autophagic flux. Through KEAP1, p62 converts this information into a transcriptional response.

However, whether this signaling node is also regulated by export from the cell is unclear. The p62 protein is detectable in human plasma and its abundance tracks disease severity in Charcot–Marie–Tooth disease type 1A (Yoon et al., 2024). This implies that a physiological secretory route exists. Moreover, p62 acts as a receptor that loads the Alzheimer-associated ubiquitin variant ubiquitin B (UBB^+1^) into autophagosome-derived secretory vesicles (Wagh and Glickman, 2025). This places p62 upstream of an unconventional secretion event. However, whether p62 itself is a secreted cargo, in what physical form it leaves the cell, which routes are responsible, or whether its export has signaling consequences has not been established. Specifically, progress has been limited by methodology. In particular, p62 is present in conditioned medium at concentrations far below the detection limit of immunoblotting, and EV purification is poorly quantitative. Furthermore, overexpression, which is the usual solution, drives artifactual p62 body formation and distorts the oligomerization behavior under study.

To address these limitations, we appended the 11-residue HiBiT peptide to endogenous p62 using Clustered Regularly Interspaced Short Palindromic Repeat (CRISPR)–CRISPR-associated protein 9 (Cas9)-mediated knock-in, so that complementation with recombinant LgBiT reconstitutes NanoLuc luciferase and quantitatively reports p62 abundance across four orders of magnitude (Dixon et al., 2016; Schwinn et al., 2018). Considering that LgBiT is membrane-impermeant, comparing the luminescence in the presence and absence of a mild detergent additionally reports whether the tagged protein resides inside a sealed membrane. Applying this system to a matched panel of knock-in lines, we found that p62 is constitutively released in membrane-enclosed carriers at a rate comparable with that of LC3B. Moreover, lysosomal inhibition redirects p62 from degradation to secretion, and autophagy-dependent and -independent routes operate in parallel. Furthermore, PB1-mediated self-oligomerization, rather than any single binding interface, supports export. In addition, KEAP1 is also exported in membrane-enclosed carriers through a RAB27-dependent route, whose disruption lowers NRF2 transcriptional output. These results identify the vesicular export of the p62–KEAP1 module as an additional input into the cellular oxidative stress response.

## Results

### Endogenous HiBiT tagging permits quantitative tracking of p62

To follow endogenous p62 without perturbing its expression, we used CRISPR-Cas9 to insert the HiBiT coding sequence immediately upstream of the stop codon of *SQSTM1* in HEK293T cells. This generated p62–HiBiT knock-in (KI) cells (Figure 1A). Starting with this line, we disrupted autophagy-related 5 (*ATG5*) or *RB1CC1* (focal adhesion kinase family interacting protein of 200 kD [FIP200]) to obtain p62–HiBiT KI/ATG5 knockout (KO) and p62–HiBiT KI/FIP200 KO cells. Sanger sequencing combined with CRISPR-ID analysis (Dehairs et al., 2016) confirmed that all edits were homozygous (Figure 1A).

**Figure 1.**
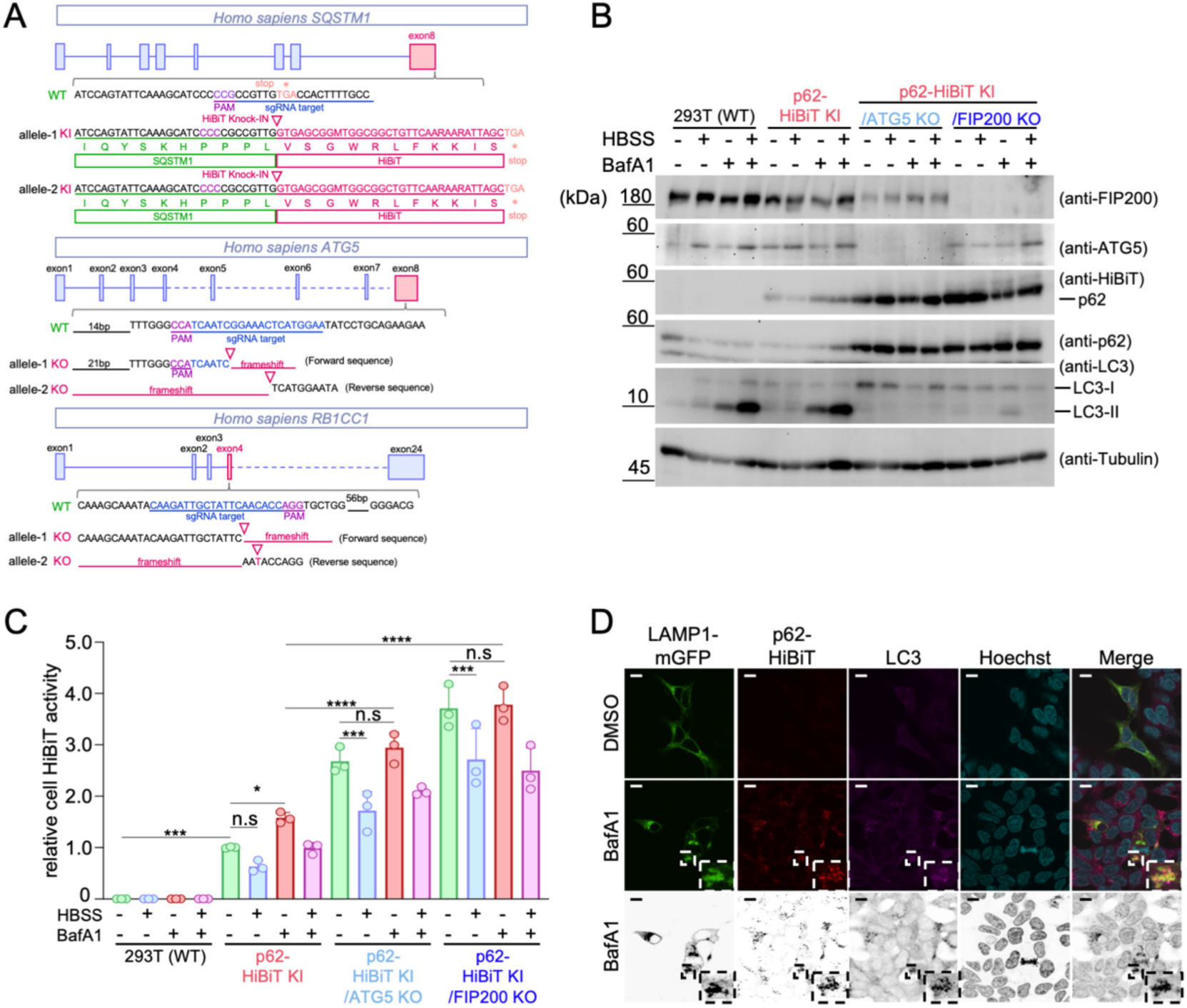
Establishment and validation of p62–HiBiT knock-in HEK293T cells. (A) Schematic of the human *SQSTM1* locus. The HiBiT insertion site is shown in red. For p62–HiBiT KI/ATG5 KO and p62–HiBiT KI/FIP200 KO cells, the edited regions resulting from gene disruption are shown in red. (B) Immunoblot analysis of p62–HiBiT and autophagy-related proteins. Wild-type (WT), p62–HiBiT KI, p62–HiBiT KI/ATG5 KO and p62–HiBiT KI/FIP200 KO HEK293T cells were treated for 6 h with or without 100 nM bafilomycin A1 (BafA1) or starved in HBSS. Lysates were probed for FIP200, ATG5, HiBiT (p62–HiBiT), p62, LC3 and α-tubulin. (C) Quantification of HiBiT-dependent NanoLuc luminescence in cell lysates. Data are presented as mean ± SD (n = 3 technical replicates); the experiment was performed independently at least twice with comparable results. Values were log-transformed. Parental WT cells (background) were excluded from the two-way ANOVAs and compared with p62–HiBiT KI cells under control conditions by lognormal Welch’s t test. The effects of BafA1 and HBSS were analyzed in separate two-way ANOVAs (genotype × BafA1; genotype × HBSS), each followed by Šídák’s multiple comparison test within each genotype; *p < 0.05, ***p < 0.001; ns, not significant. (D) BafA1 induces lysosomal accumulation of p62–HiBiT. The p62–HiBiT KI cells transiently expressing LAMP1–mGFP (green) were treated with 100 nM BafA1 for 6 h or left untreated, fixed, and stained for HiBiT (red), LC3 (magenta) and with Hoechst 33342 (cyan). Lower panels show binarized images of extracted puncta generated in Fiji. Insets show magnified views of the boxed regions. Scale bar, 10 μm. Abbreviations: SQSTM1, sequestosome 1; KI, knock-in; ATG5, autophagy-related 5; KO, knockout; FIP200, focal adhesion kinase family interacting protein of 200 kD; HBSS, Hanks’ Balanced Salt Solution; LC3, microtubule-associated protein 1 light chain 3; SD, standard deviation; ANOVA, analysis of variance; LAMP1, lysosomal-associated membrane protein 1; mGFP, membrane-targeted green fluorescent protein.

Immunoblotting with an anti-HiBiT antibody detected a single species that co-migrated with p62 with no additional bands. This confirmed that the tag was expressed exclusively as a p62 fusion (Figure 1B). LC3-II accumulated to a similar extent in p62–HiBiT KI and parental wild-type (WT) cells under starvation and under Bafilomycin A1 (BafA1)-induced lysosomal inhibition. This indicated that autophagy induction is unimpaired in the KI line. As expected, LC3-II formation was undetectable in ATG5 KO cells and was strongly reduced in FIP200 KO cells. Both lines accumulated p62, thereby confirming a functional block in the autophagy pathway (Figure 1B).

Luminescence measured after LgBiT complementation gave a robust signal in p62–HiBiT KI lysates and only a background signal in parental cells (Figure 1C). In p62–HiBiT KI cells, starvation in Hanks’ Balanced Salt Solution (HBSS) did not significantly change the intracellular HiBiT activity, whereas BafA1 increased it significantly. This BafA1-induced increase was absent in ATG5 KO and FIP200 KO cells, which indicates that the accumulating pool was normally cleared by the autophagy–lysosome pathway (Figure 1C).

Immunofluorescence showed the same pattern at the localization level. Under steady-state conditions, p62–HiBiT was distributed as small, dispersed cytoplasmic puncta. After BafA1 treatment, it was redistributed into prominent clusters that co-localized with lysosomal-associated membrane protein 1 (LAMP1) and LC3 (Figure 1D). This behavior was consistent with that reported for untagged endogenous p62 (Bjørkøy et al., 2005), thus indicating that the HiBiT tag does not alter p62 localization or trafficking. Therefore, the KI system faithfully reports endogenous p62 dynamics and is suitable for quantifying its extracellular release.

### The p62 protein is constitutively released in membrane-enclosed carriers

Next, we investigated whether p62 reaches the extracellular space. The conditioned medium was cleared of cells and debris and separated by differential centrifugation into a 10,000 × *g* pellet (10k ppt; enriched in large EVs), a 100,000 × *g* pellet (100k ppt; enriched in small EVs), and the residual non-sedimentable supernatant (100k sup), following the EV nomenclature and reporting recommendations (Welsh et al., 2024) (see Materials and methods). The same fractionation scheme, applied to the same cell lines and culture conditions, has been characterized previously (Maeda et al., 2024; Saito et al., 2026). Specifically, the 10k and 100k pellets are positive for the transmembrane markers cluster of differentiation (CD)9, CD63, and CD81 and for the cytosolic markers apoptosis-linked gene 2-interacting protein X (ALIX) and tumor susceptibility gene 101 (TSG101), are depleted of the non-EV markers calnexin and GM130, and contain particles of the expected size distribution and abundance by nanoparticle tracking analysis. In contrast, the 100k supernatant is largely devoid of particles and of these markers. Therefore, we refer to the pellet fractions as membrane-enclosed carriers.

To quantify p62 release, we compared it with two reference proteins tagged in the same way: translocase of outer mitochondrial membrane 20 (TOMM20)–HiBiT, which is an outer mitochondrial membrane protein, and HiBiT–LC3B, which is an autophagosome marker (Saito et al., 2026) (Figure 2A). The intracellular abundance of p62–HiBiT at steady state was lower than that of either reference protein (Figure 2B). Nevertheless, when the HiBiT signal recovered from each extracellular fraction was expressed relative to the corresponding total cellular plus extracellular signal, the secretion rate of p62–HiBiT was comparable with that of HiBiT–LC3B in both the 10k ppt and 100k ppt fractions, and the majority of secreted p62 was recovered in the 10k ppt fraction. The secretion rate of p62 was lower than that of TOMM20–HiBiT (Figure 2C).

**Figure 2.**
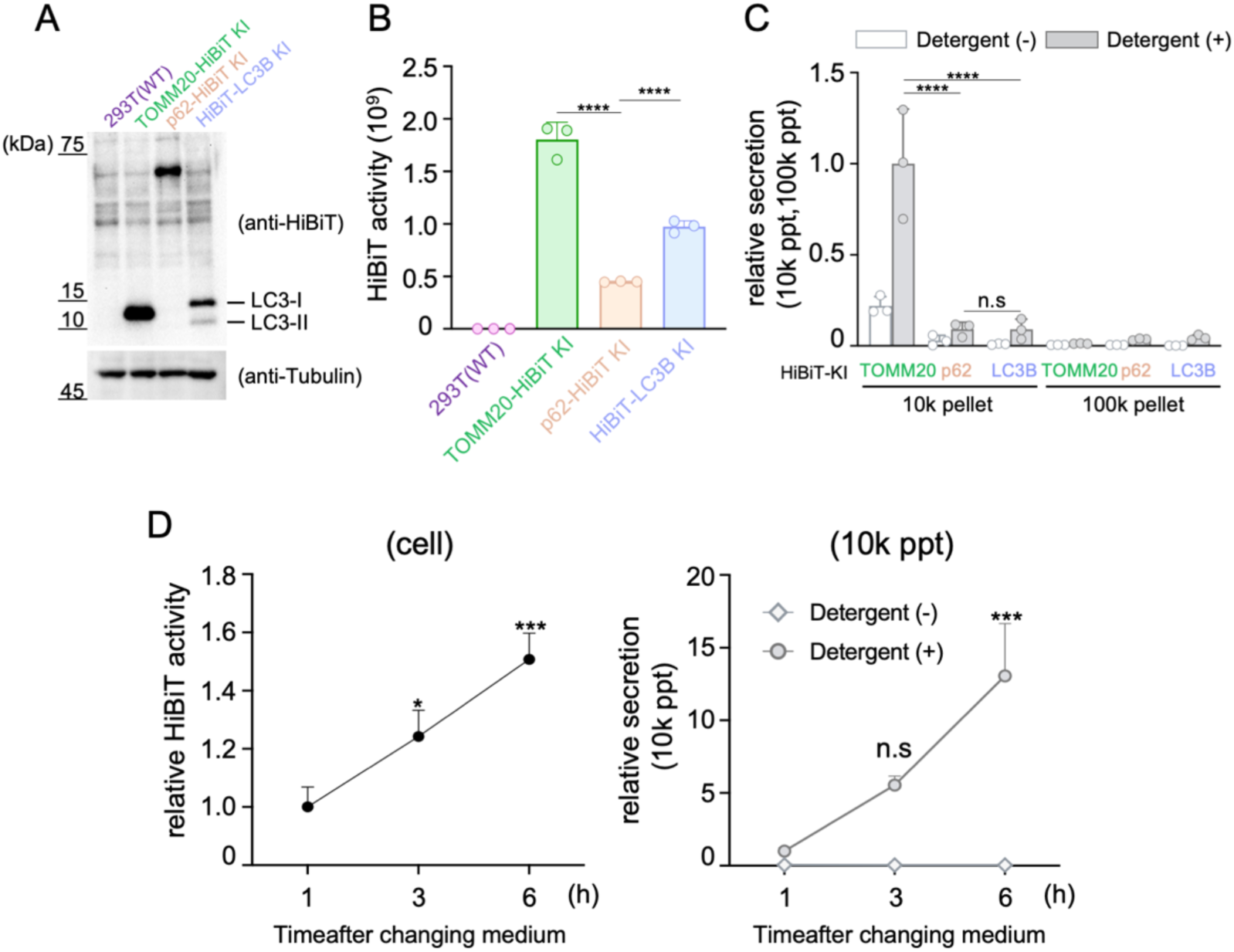
Secreted p62–HiBiT is recovered in membrane-enclosed extracellular fractions. (A) Immunoblot confirming expression of the HiBiT fusion proteins. Lysates of WT, TOMM20–HiBiT KI, p62–HiBiT KI and HiBiT–LC3B KI HEK293T cells were probed for TOMM20, p62, LC3B, and α-tubulin, and with an anti-HiBiT antibody (upper); α-tubulin is shown as loading control (lower). (B) Comparison of expression levels of the HiBiT fusion proteins. Cells were cultured for 48 h, and HiBiT-dependent NanoLuc luminescence was measured in lysates. Values were log-transformed, and parental WT cells (background) were excluded from the analysis. One-way ANOVA with Dunnett’s multiple comparison test against p62–HiBiT KI; ****p < 0.0001. (C) Secretion rate and detergent dependence. Conditioned medium collected after 48 h was fractionated into the 10,000 × *g* pellet (10k ppt; left) and the 100,000 × *g* pellet (100k ppt; right), and HiBiT activity was measured with or without 0.1% Triton X-100. The secretion rate was calculated as the HiBiT activity in the indicated extracellular fraction divided by the sum of the activity in that fraction and in the corresponding cell fraction. Data are presented as mean ± SD (n = 3 technical replicates); the experiment was performed independently at least twice with comparable results. Two-way ANOVA with Tukey’s multiple comparison test; ****p < 0.0001; ns, not significant. (D) Time-dependent accumulation and release of p62–HiBiT. HiBiT activity was measured in cell lysates (left) and conditioned medium (right) at the indicated times after a medium change; medium was measured with and without 0.1% Triton X-100. Data are presented as mean ± SD (n = 3). One-way ANOVA with Dunnett’s multiple comparison test; *p < 0.05, ***p < 0.001; ns, not significant. Abbreviations: WT, wild type; TOMM20, translocase of outer mitochondrial membrane 20; KI, knock-in; LC3B, microtubule-associated protein 1 light chain 3 beta; ANOVA, analysis of variance; SD, standard deviation.

Considering that LgBiT cannot cross an intact lipid bilayer, luminescence measured without detergent reports only the protein exposed to the buffer, whereas the signal obtained after the addition of 0.1% Triton X-100 reports the total protein. For p62–HiBiT, HiBiT–LC3B, and TOMM20–HiBiT, extracellular signals in both pellet fractions were detected almost exclusively in the presence of detergent (Figure 2C). This indicated that the secreted p62 was enclosed within a sealed membrane.

The release was progressive rather than burst-like. After exchange with fresh medium, HiBiT activity increased with time in both the cell lysate and secreted fraction, and the extracellular signal remained detergent-dependent (Figure 2D). Therefore, we conclude that p62 is constitutively exported in membrane-enclosed carriers at a fractional rate similar to that of LC3B but substantially lower than that of TOMM20.

### Lysosomal inhibition redirects p62 from degradation to secretion

Secretory autophagy is activated when lysosomal degradation is compromised (Solvik et al., 2022). Treatment of p62–HiBiT KI cells with BafA1 for 6 h caused the expected intracellular accumulation of p62–HiBiT (Figure 3A). In parallel, this treatment increased the secretion rate of the 10k ppt and 100k ppt fractions, with a larger relative increase in the 100k ppt fraction (Figure 3B).

**Figure 3.**
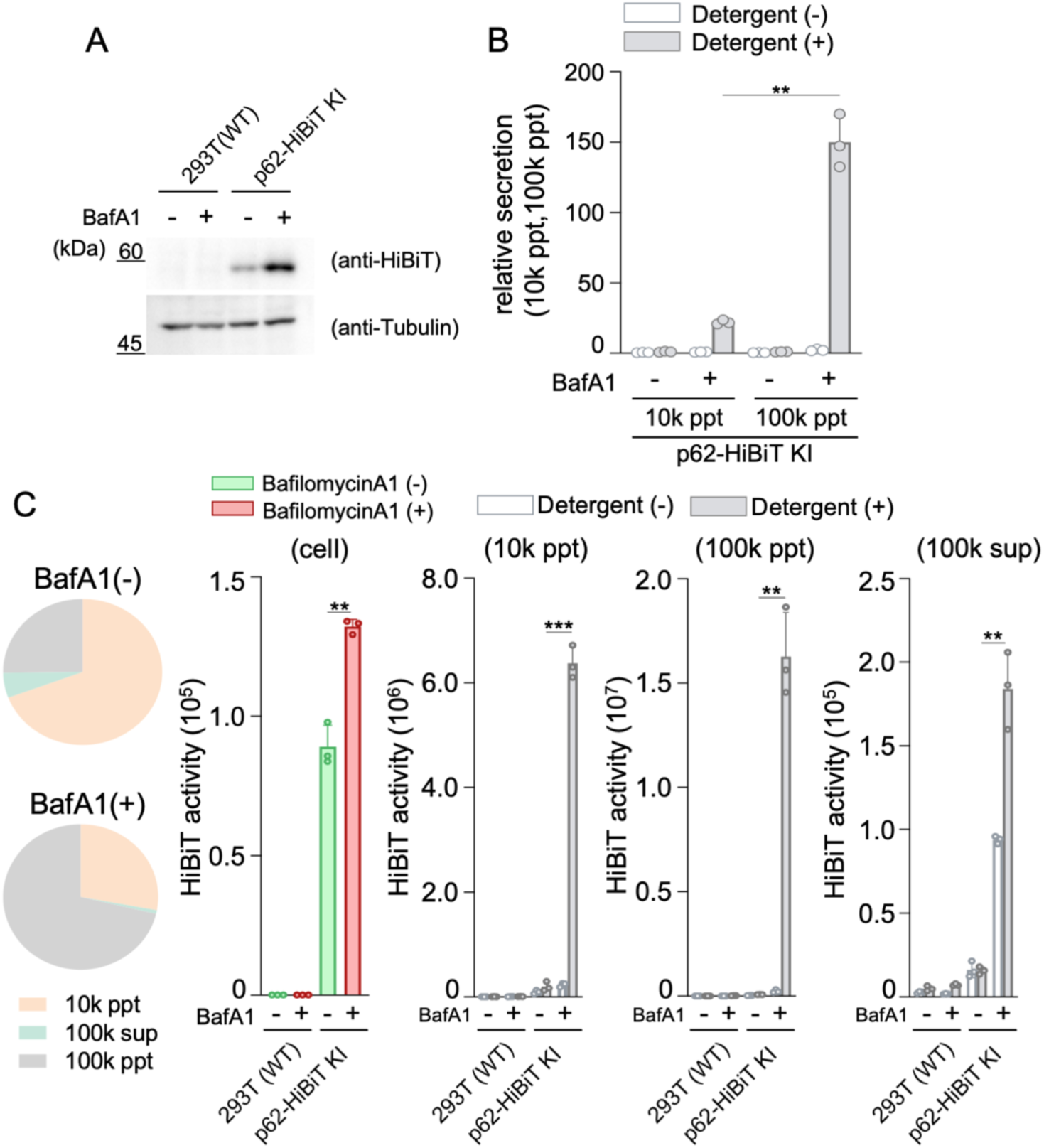
Lysosomal inhibition increases p62–HiBiT release. (A) Intracellular p62–HiBiT after BafA1 treatment. WT and p62–HiBiT KI HEK293T cells were treated with or without 100 nM BafA1 for 6 h, and lysates were probed with anti-HiBiT (upper) and anti-α-tubulin (lower). (B) Effect of BafA1 on p62–HiBiT secretion. The 10k ppt and 100k ppt fractions were recovered from conditioned medium of p62–HiBiT KI cells, and HiBiT activity was measured with and without 0.1% Triton X-100. Secretion rates were calculated as in Figure 2C. Data are presented as mean ± SD (n = 3 technical replicates); the experiment was performed independently at least twice. Unpaired two-tailed Student’s t test; **p < 0.01. (C) Extracellular distribution of secreted p62–HiBiT. The p62–HiBiT KI cells treated with or without 100 nM BafA1 for 6 h were separated via differential centrifugation into cell, 10k ppt, 100k ppt, and 100k sup fractions, and HiBiT activity in each fraction was measured with and without 0.1% Triton X-100. Distributions are shown as pie charts. Data are presented as mean ± SD (n = 3); the experiment was performed independently at least twice. Unpaired two-tailed Student’s t test; **p < 0.01, ***p < 0.001. Abbreviations: BafA1, bafilomycin A1; WT, wild type; KI, knock-in; SD, standard deviation.

To determine how the secreted material was distributed among these fractions, cells cultured for 48 h were switched to BafA1-containing medium for an additional 6 h, and the cells and conditioned medium were resolved into cell, 10k ppt, 100k ppt, and 100k sup fractions (Figure 3C). BafA1 increased p62–HiBiT in all extracellular fractions relative to those in the untreated control. In the 10k ppt and 100k ppt fractions, the signal was detergent-dependent, both with and without BafA1. This confirmed that the membrane-enclosed form dominated and lysosomal inhibition did not change the physical form of this pool.

The 100k sup fraction behaved differently. Here, a substantial part of the p62–HiBiT signal was detected in the absence of detergent, both at steady state and after BafA1 treatment (Figure 3C). Therefore, a fraction of extracellular p62 is not membrane-enclosed. This indicates that p62 leaves the cell in at least two physically distinct forms, and these forms are unlikely to arise from a single carrier.

### Autophagy-dependent and -independent components of p62 release

To separate these components genetically, we compared isogenic p62–HiBiT KI, p62–HiBiT KI/ATG5 KO, and p62–HiBiT KI/FIP200 KO lines. In p62–HiBiT KI cells, BafA1 markedly increased the secretion rate, whereas starvation in HBSS had little effect relative to the steady state. When the two treatments were combined, the BafA1-driven increase was substantially attenuated (Figure 4A). Thus, p62 release is governed by the state of the degradation pathway rather than by autophagy induction.

**Figure 4.**
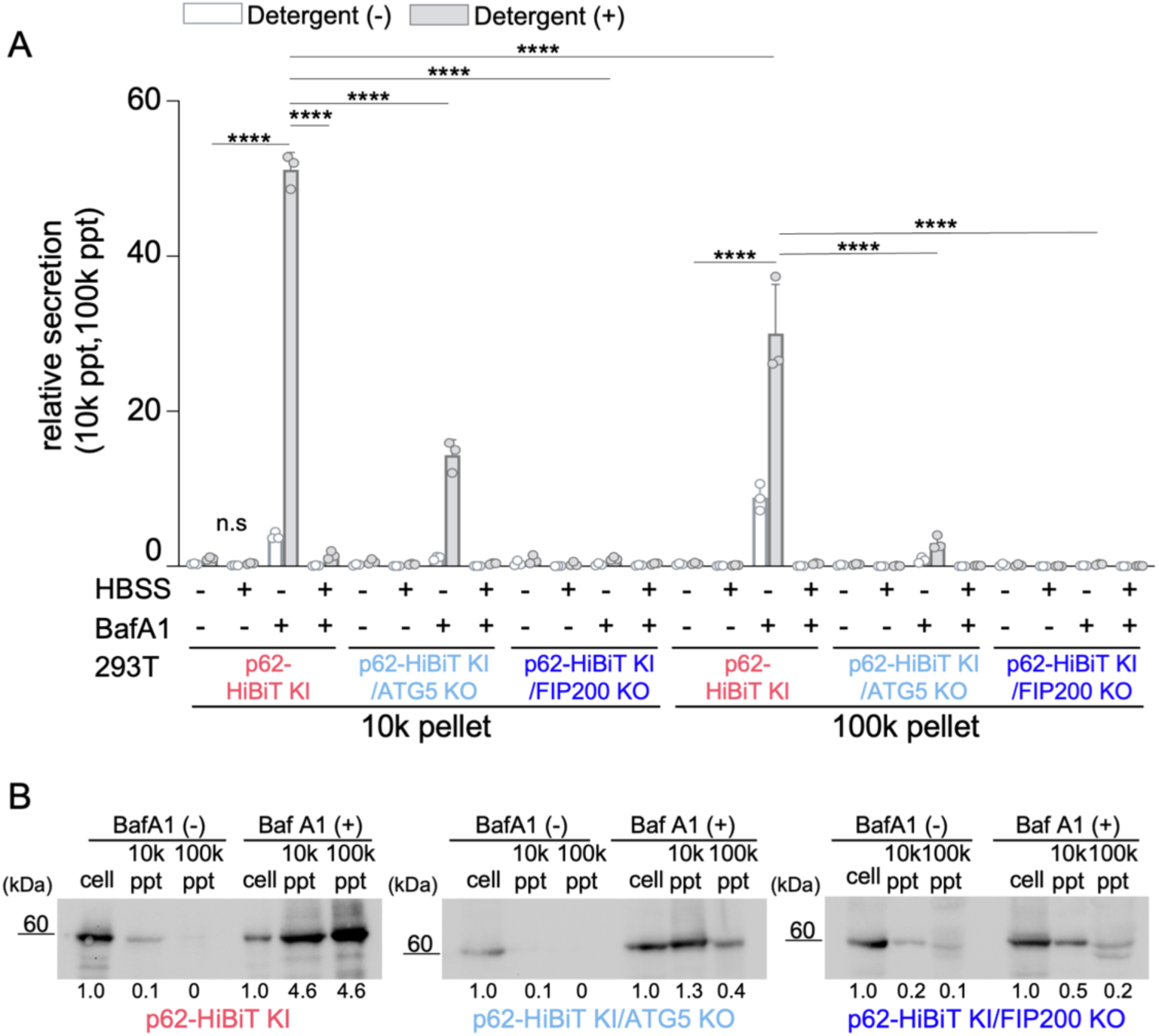
The BafA1-inducible component of p62–HiBiT release requires ATG5 and FIP200. (A) Effect of autophagy gene disruption on p62–HiBiT secretion. The p62–HiBiT KI, p62–HiBiT KI/ATG5 KO and p62–HiBiT KI/FIP200 KO cells were cultured for 6 h under control conditions, with 100 nM BafA1, in HBSS, or in HBSS containing 100 nM BafA1. Cells and conditioned medium were collected, and HiBiT activity was measured with or without 0.1% Triton X-100. Secretion rates were calculated from the 10k ppt (left) and 100k ppt (right) fractions as in Figure 2C. Data are presented as mean ± SD (n = 3 technical replicates); the experiment was performed independently at least twice. Two-way ANOVA with Tukey’s multiple comparison test; ****p < 0.0001; ns, not significant. (B) Immunoblot detection of secreted p62–HiBiT. Cells of the indicated genotypes were cultured for 12 h with or without 100 nM BafA1. Conditioned medium was fractionated via differential centrifugation, and p62–HiBiT in each fraction was detected with an anti-HiBiT antibody. Numbers below each panel indicate the amount of p62–HiBiT in the 10k ppt or 100k ppt fraction relative to the cell fraction, based on band intensity. Abbreviations: KI, knock-in; ATG5, autophagy-related 5; KO, knockout; BafA1, bafilomycin A1; HBSS, Hanks’ Balanced Salt Solution; SD, standard deviation; ANOVA, analysis of variance.

In ATG5 KO cells, the steady-state secretion rate was indistinguishable from that of the parental KI line. However, the BafA1-dependent increase was strongly reduced. FIP200 KO cells behaved similarly (Figure 4A). Therefore, the released component of p62 that is mobilized by lysosomal inhibition depends on phagophore initiation and LC3 lipidation, whereas the constitutive component does not.

Immunoblotting of the secreted fractions yielded similar results at the protein level (Figure 4B). After 12 h in the presence of BafA1, p62–HiBiT was readily detected in the 10k ppt and 100k ppt fractions of KI cells, and this signal was reduced in ATG5 KO and FIP200 KO cells. At steady state, secreted p62–HiBiT remained below the detection limit of immunoblotting in all lines. This emphasizes the sensitivity of the complementation assay.

Collectively, these data define two components of p62 release: a constitutive, ATG5/FIP200-independent component and an inducible component that is engaged when lysosomal degradation is blocked. Only the inducible component requires the core autophagy machinery. The same two-component behavior has been described for LC3B (Saito et al., 2026), which suggests that p62 and LC3B use partially overlapping routes.

### PB1-mediated self-oligomerization, but not LC3 or KEAP1 binding, supports p62 secretion

Structure–function analysis of p62 requires the expression of mutant proteins, but conventional cytomegalovirus (CMV)-driven overexpression produces supraphysiological p62 concentrations that force p62 body assembly and would confound any measurement of oligomerization-dependent secretion. Therefore, we generated a series of vectors carrying progressive 5′ deletions of the CMV promoter (CMVΔ1.0 to CMVΔ5.0) (Morita et al., 2012) driving FLAG–HiBiT-tagged p62 (Figure 5A). Expression decreased stepwise with the extent of promoter deletion (Figure 5B), with a corresponding reduction in the number and size of p62 puncta relative to the full-length CMV promoter construct (Figure 5C). The most extensively truncated construct, CMVΔ5.0, yielded intracellular HiBiT activity 6.52-fold that of endogenous p62–HiBiT in knock-in cells without BafA1 and 4.93-fold with BafA1 (Figures 5D–5F). Functionally, p62 expressed from CMVΔ5.0 was released into the 10k ppt fraction, where HiBiT activity measured in the presence of detergent was 1.27-fold that of knock-in cells without BafA1 and 1.19-fold with BafA1. As observed for endogenous p62, the extracellular HiBiT activity was detergent dependent under all conditions (Figures 5G and 5H). CMVΔ5.0 therefore provides a low-level expression platform, within an order of magnitude of endogenous p62, suitable for domain analysis.

**Figure 5.**
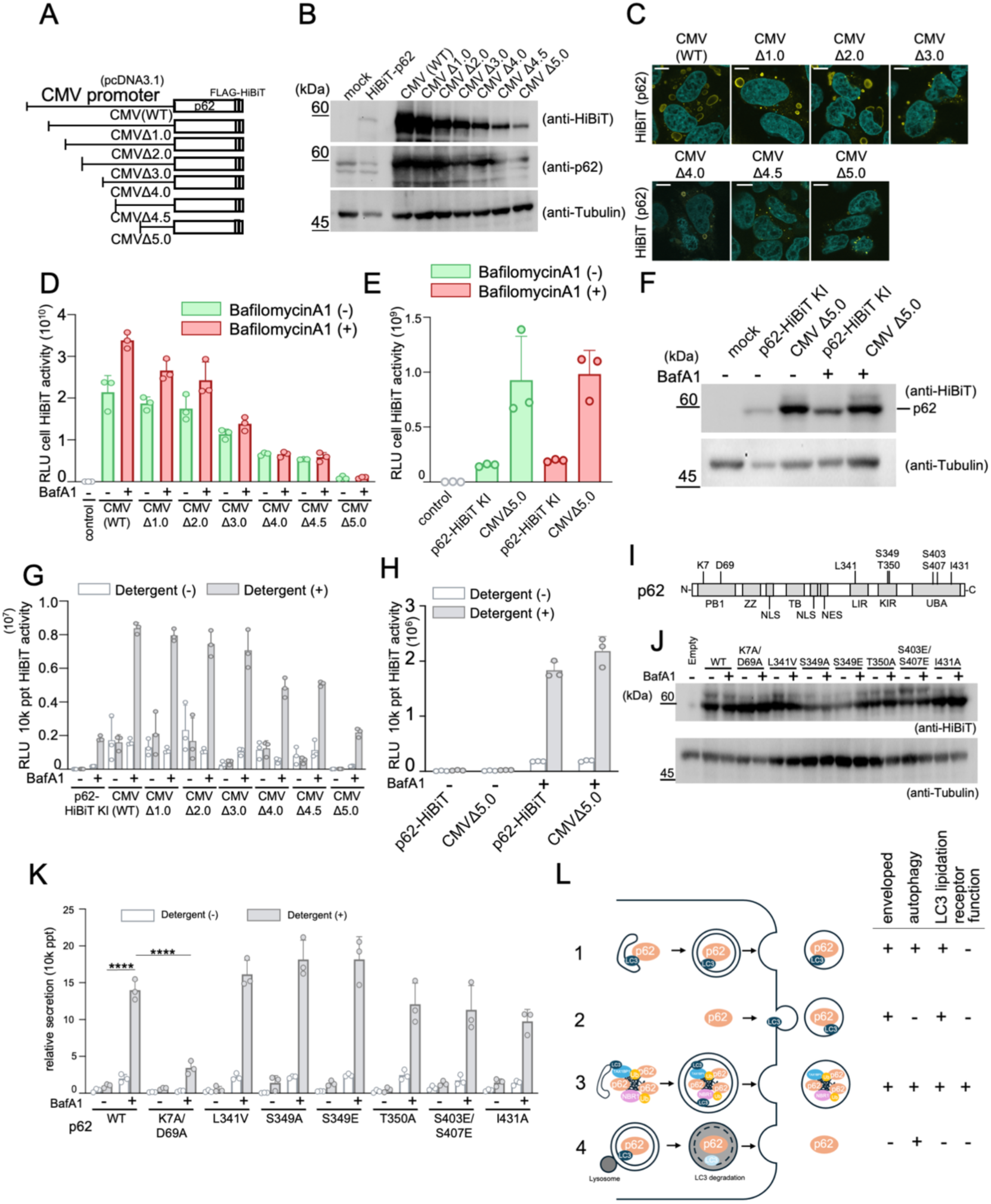
An attenuated CMV promoter lowers p62 expression toward endogenous levels and reveals that PB1-mediated self-oligomerization supports p62 secretion. (A) Schematic of the p62 expression vectors. FLAG–HiBiT-tagged p62 was placed under the control of the full-length CMV promoter or a series of 5′ deletion variants (CMVΔ1.0–CMVΔ5.0). (B) Comparison of p62 expression from each construct. WT HEK293T cells were transfected with the indicated plasmids, and lysates prepared after 24 h were probed for FLAG–HiBiT–p62 (upper), endogenous p62 (middle), and α-tubulin (lower) with the respective antibodies or with an anti-HiBiT antibody. (C) Localization of transiently expressed p62. WT HeLa cells transfected with the indicated plasmids were fixed after 24 h and stained for anti-HiBiT antibody (yellow) and with Hoechst 33342 (cyan). Scale bar, 10 μm. (D) Intracellular HiBiT activity of transiently expressed p62. WT HEK293T cells transfected with the indicated plasmids were treated after 24 h with or without 100 nM bafilomycin A1 (BafA1) for 6 h, and HiBiT activity was measured in lysates. Data are presented as mean ± SD (n = 3 technical replicates). (E) Expanded view of (D) comparing p62–HiBiT KI cells with cells expressing the CMVΔ5.0 construct. No statistical test was applied; fold differences are given in the main text. (F) Immunoblot validation of p62 expression. Lysates obtained under the conditions in (D) were probed for FLAG–HiBiT–p62 (upper) and α-tubulin (lower). (G) Detection of secreted p62–HiBiT. Conditioned medium from the experiment in (D) was collected and HiBiT activity was measured with or without 0.1% Triton X-100. Data are mean ± SD (n = 3). (H) Expanded view of (G) comparing p62–HiBiT KI cells with cells expressing the CMVΔ5.0 construct. No statistical test was applied. (I) Domain organization of p62, showing the known functional domains. PB1: Phox/Bem1p domain, ZZ: Zinc finger domain, NLS: nuclear localization sequences, NES: nuclear export sequence, TB: TRAF binding domain, LIR: LC3-interacting region, KIR: Keap1-interacting region, UBA: Ubiquitin-associated domain. (J) Expression of the p62 variants. WT HEK293T cells were transfected with the indicated HiBiT-tagged p62 constructs (wild type and the individual mutants), and lysates prepared after 24 h were probed for FLAG–HiBiT–p62 (upper) and α-tubulin (lower) with the respective antibodies or with an anti-HiBiT antibody. (K) Secretion rates of the p62 variants. WT HEK293T cells transiently expressing each HiBiT-tagged p62 were treated after 24 h with or without 100 nM BafA1 for 6 h. Cells and conditioned medium were collected, and HiBiT activity was measured with or without 0.1% Triton X-100. Secretion rates were calculated as the HiBiT activity in the 10k ppt fraction divided by the sum of the activity in that fraction and in the cell fraction, as in Figure 2C. Data are presented as mean ± SD (n = 3 technical replicates); the experiment was performed independently at least twice. Two-way ANOVA with Tukey’s multiple comparison test; ****p < 0.0001. (L) Model for p62 secretion pathway. The p62 may be released through multiple routes: (1) an autophagy-dependent route in which LC3-bound p62 is incorporated into autophagosomes and secreted; (2) an autophagy-independent route that nonetheless involves incorporation into membrane vesicles; (3) secretion as a higher-order complex formed through interaction with other receptor proteins; and (4) release of non-enveloped p62 following lysosomal degradation. Abbreviations: CMV, cytomegalovirus; WT, wild type; SD, standard deviation; KI, knock-in; ANOVA, analysis of variance; LC3B, microtubule-associated protein 1 light chain 3 beta.

Using this platform, we expressed a panel of HiBiT-tagged p62 variants targeting each known functional interface (Figure 5I): K7A/D69A (PB1 oligomerization-deficient), L341V (an ALS/FTLD-associated LIR variant with impaired LC3B binding (Goode et al., 2016)), S349A and S349E (KIR phospho-null and phosphomimetic (Ichimura et al., 2013)), T350A, S403E/S407E (UBA phosphomimetic (Matsumoto et al., 2011)) and I431A (UBA). All the variants were expressed at comparable levels (Figure 5J).

Every variant retained the qualitative behavior of wild-type p62. In particular, BafA1 increased the secretion rate, and extracellular HiBiT activity was detergent-dependent. This indicated that none of these mutations changed the membrane-enclosed nature of the secreted species (Figure 5K). However, quantitatively, the PB1 mutant K7A/D69A was distinct. Specifically, its BafA1-dependent secretion was substantially lower than that of WT p62 (Figure 5K). Considering that this mutant failed to form p62 bodies (Itakura and Mizushima, 2011), this result indicated that self-oligomerization — that is, the formation of a higher-order p62 assembly — is a prerequisite for efficient export. In contrast, the LIR variant L341V showed no considerable change in BafA1-dependent secretion (Figure 5K). Therefore, direct LC3 engagement by p62 was dispensable. A plausible explanation is that other receptors present in p62 bodies, such as neighbor of BRCA1 gene 1 (NBR1) (Kirkin et al., 2009) and Tax1-binding protein 1 (TAX1BP1) (Turco et al., 2021), provide contact with LC3 in *trans* to ensure that an oligomer is captured by the phagophore, even when the LIR of p62 is compromised. Neither the KIR nor UBA mutations altered secretion (Figure 5K).

These properties of membrane enclosure, dependence on the lysosomal status, partial dependence on the core autophagy machinery, and independence of a single LC3-binding interface are summarized in Figure 5L. This frames four non-exclusive routes: an autophagy-dependent route, an autophagy-independent route that yields membrane-enclosed p62, secretion as part of a higher-order receptor assembly, and release of non-enveloped p62. Each of these is considered in the Discussion.

### KEAP1, like p62, is secreted in membrane-enclosed carriers

Under conditions of insufficient autophagic flux, the accumulation of p62 sequesters KEAP1, thereby stabilizing NRF2 (Ichimura et al., 2013; Komatsu et al., 2010; Yamamoto et al., 2018). If p62 is exported, KEAP1 may also be exported with it. Therefore, we determined whether KEAP1 was secreted and whether its secretion depended on p62 binding.

We generated HiBiT-tagged KEAP1 carrying a single (R380A, R415A, and R483A) or combined substitution in the Kelch domain pocket that mediates binding to the ETGE/STGE motifs (Padmanabhan et al., 2006) of NRF2 and p62 (Figures 6A and 6B). Affinity purification of One-Strep–FLAG–p62 confirmed that these substitutions reduced the association between KEAP1 and p62 (Figure 6C).

**Figure 6.**
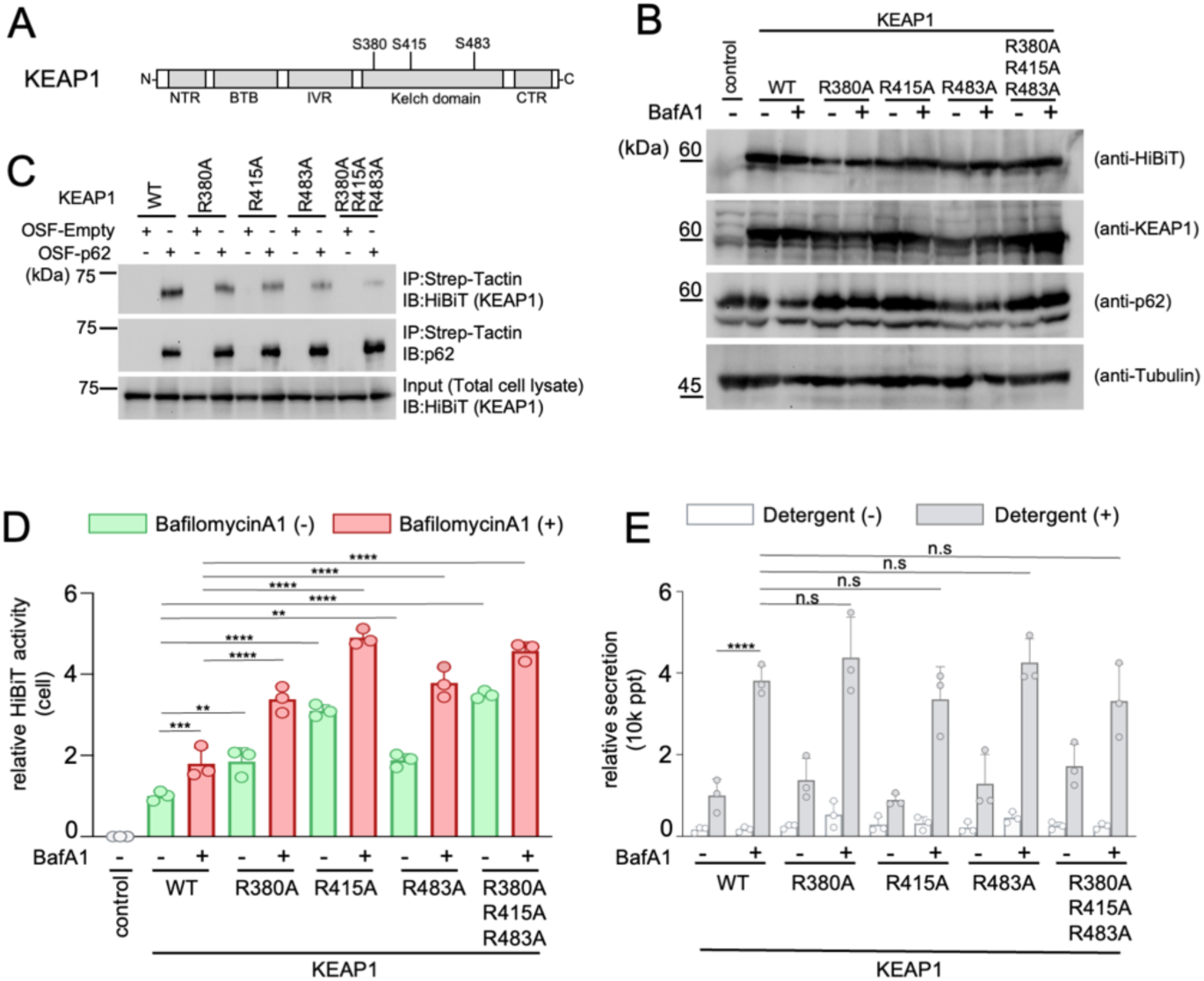
KEAP1 is secreted in membrane-enclosed carriers independently of p62 binding. (A) Domain organization of KEAP1 with the positions of the mutated Kelch-domain residues. NTA: N-terminal region, BTB: Broad complex, Tramtrack, and Bric-à-Brac domain, IVR: Intervening Region, CTR: C-terminal region. (B) Expression of the KEAP1 variants. WT HEK293T cells were transfected with the indicated HiBiT-tagged KEAP1 constructs, and lysates prepared after 24 h were probed for FLAG–HiBiT–KEAP1 (first panel), endogenous KEAP1 (second), endogenous p62 (third), and α-tubulin (fourth). (C) KEAP1–p62 interaction. Each HiBiT–KEAP1 construct was co-expressed with One-Strep–FLAG (OSF)–p62 in HEK293T cells and affinity purified on Strep-Tactin resin. Purified fractions were immunoblotted for HiBiT–KEAP1 (upper) and p62 (middle) and compared with HiBiT–KEAP1 in the input total cell lysate (lower). (D) Intracellular accumulation of KEAP1. WT HEK293T cells transiently expressing each HiBiT–KEAP1 were treated after 24 h with or without 100 nM BafA1 for 6 h, and HiBiT activity was measured in lysates. Data are presented as mean ± SD (n = 3 technical replicates); the experiment was performed independently at least twice. Two-way ANOVA with Tukey’s multiple comparison test; **p < 0.01, ***p < 0.001, ****p < 0.0001. (E) Secretion rates of the KEAP1 variants, determined from the 10k ppt fraction as in Figure 2C under the conditions in (D). Data are presented as mean ± SD (n = 3 technical replicates); the experiment was performed independently at least twice. Two-way ANOVA with Tukey’s multiple comparison test; ****p < 0.0001; ns, not significant. Abbreviations: KEAP1, Kelch-like ECH-associated protein 1; WT, wild type; BafA1, bafilomycin A1; SD, standard deviation; ANOVA, analysis of variance.

BafA1 caused the intracellular accumulation of both WT and mutant KEAP1, and this accumulation was more pronounced for the mutants than for the WT protein, irrespective of treatment (Figure 6D). This was consistent with the binding-dependent turnover of KEAP1. Secretion rates increased upon BafA1 treatment for WT KEAP1 and all mutants, and extracellular HiBiT activity was detergent-dependent (Figure 6E). Therefore, KEAP1 was exported in membrane-enclosed carriers in the same manner as p62 and LC3B.

Notably, the BafA1-dependent secretion rate did not differ substantially between the WT KEAP1 and binding-deficient mutants (Figure 6E). Under lysosomal blockade, KEAP1 export does not require its interaction with p62. Specifically, the two proteins appear to reach the same carriers via routes that can operate independently. One plausible explanation is that KEAP1 is recruited to the site of capture as a client of p62 condensates rather than through the KIR–Kelch interface, a point we return to in the Discussion.

### RAB27-dependent export of the p62–KEAP1 module is required for full NRF2 transcriptional output

RAB27 GTPases control the docking and fusion of multivesicular bodies with the plasma membrane and are required for EV release (Ostrowski et al., 2010). To test whether the p62–KEAP1 module uses this route, we co-expressed WT RAB27A or the dominant-negative variants T23N and N133I together with HiBiT-tagged LC3B, p62, or KEAP1. Thereafter, we measured their release into the 10k ppt fraction. Both dominant-negative variants reduced the secretion of all three proteins (Figure 7A), and immunoblotting of the corresponding lysates revealed reciprocal intracellular accumulation (Figure 7B). Therefore, LC3B, p62, and KEAP1 share a RAB27-dependent export step.

**Figure 7.**
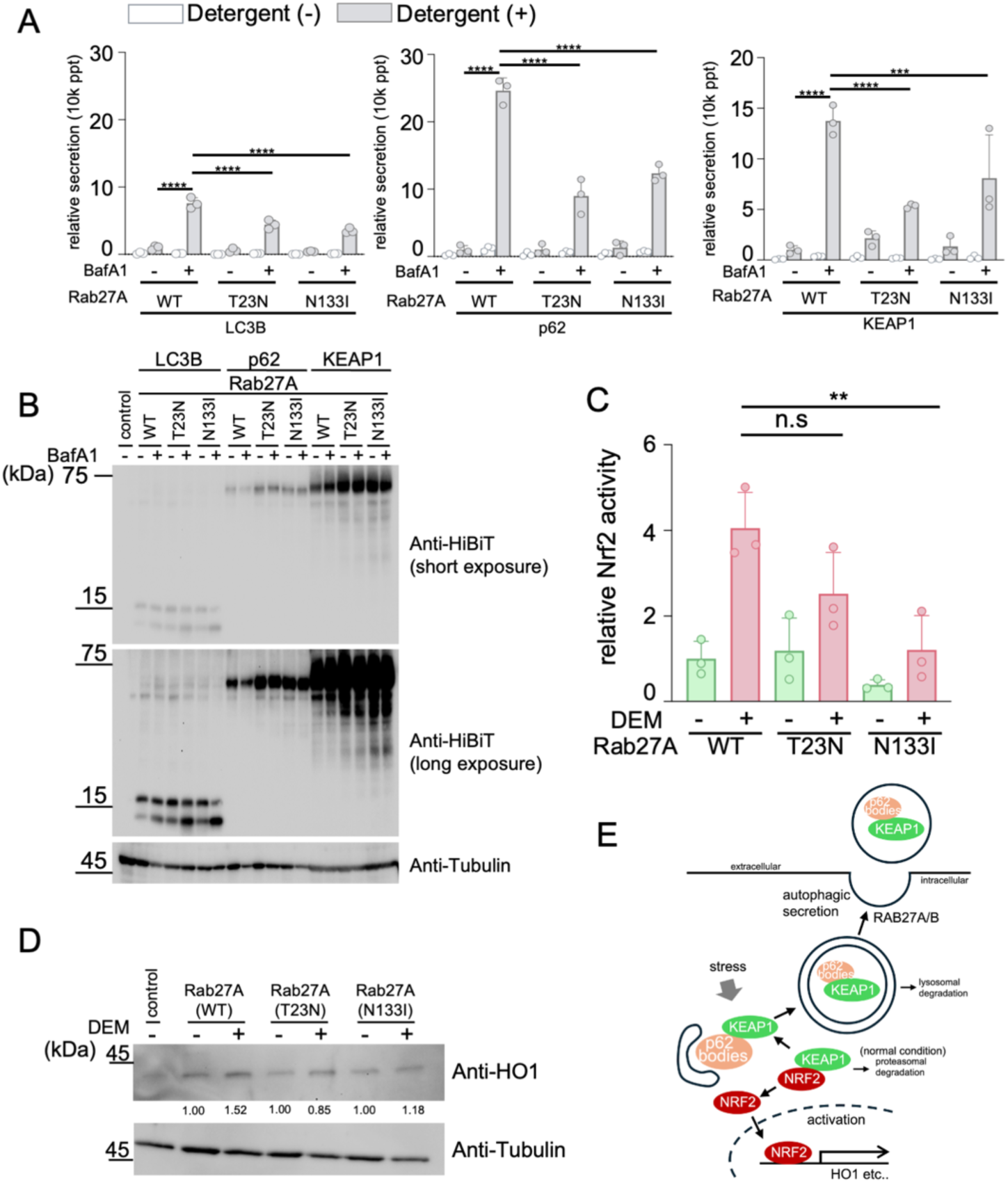
RAB27-dependent secretion of LC3B, p62, and KEAP1 is required for full NRF2 transcriptional output. (A) RAB27-dependent secretion. WT HEK293T cells were cotransfected with RAB27A WT or the dominant-negative variants T23N or N133I together with HiBiT-tagged LC3B (left), p62 (middle), or KEAP1 (right). Cells were treated with or without BafA1, and HiBiT activity in the 10k ppt fraction was measured in the presence or absence of 0.1% Triton X-100. Secretion rates were calculated as in Figure 2C. Data are presented as mean ± SD (n = 3 technical replicates); the experiment was performed independently at least twice. Two-way ANOVA with Tukey’s multiple comparison test; ***p < 0.001, ****p < 0.0001. (B) Intracellular protein levels. Lysates from the experiment in (A) were immunoblotted with anti-HiBiT at short (upper) and long (middle) exposure, and with anti-α-tubulin (lower). (C) NRF2 activity. SH-SY5Y cells were cotransfected with RAB27A WT or the dominant-negative variants together with the ARE-driven firefly luciferase reporter and the NanoLuc internal control, treated with or without 500 nM diethyl maleate (DEM) for 6 h, and assayed for luciferase activity. NRF2 activity is expressed as firefly luminescence normalized to NanoLuc luminescence. Data are presented as mean ± SD (n = 3 technical replicates); the experiment was performed independently at least twice. One-way ANOVA with Dunnett’s multiple-comparisons test; **p < 0.01; ns, not significant. (D) HO-1 expression. Lysates from the experiment in (C) were immunoblotted for HO-1 (upper) and α-tubulin (lower). DEM-induced HO-1 accumulation was reduced by both dominant-negative variants. Quantification based on band intensity is expressed relative to the RAB27A WT, DEM-untreated condition, set to 1. (E) Model for the regulation of NRF2 by the secretion of KEAP1, p62, and LC3B. In this model, RAB27A-dependent vesicular export removes KEAP1 from the cytoplasm in parallel with p62-mediated sequestration; blocking export retains KEAP1 intracellularly and is associated with a dampened NRF2 response to oxidative stress. The model is consistent with, but not established by, the present data. Abbreviations: LC3B, microtubule-associated protein 1 light chain 3 beta; KEAP1, Kelch-like ECH-associated protein 1; NRF2, nuclear factor erythroid 2-related factor 2; WT, wild type; BafA1, bafilomycin A1; SD, standard deviation; ANOVA, analysis of variance; ARE, antioxidant response element; HO-1, heme oxygenase 1.

We then investigated whether blocking this step had a signaling effect. SH-SY5Y cells expressing WT RAB27A or the dominant-negative variants were treated with diethyl maleate (DEM) to activate NRF2, and the NRF2 transcriptional output was measured using an antioxidant response element-driven luciferase reporter. Expression of either dominant-negative variant lowered the DEM-induced NRF2 activity relative to that of WT RAB27A (Figure 7C). Immunoblotting for the endogenous NRF2 target heme oxygenase 1 (HO-1) in the same lysates yielded a concordant result. In particular, DEM induced HO-1 in control cells, and this induction was reduced by the expression of the RAB27A dominant-negative variant (Figure 7D). Reporter activity and the endogenous target protein therefore moved in the same direction, indicating that the loss of RAB27-dependent export is associated with an attenuated NRF2 response at the level of target gene output.

These observations are consistent with a model (Figure 7E) in which RAB27-dependent vesicular export continuously removes KEAP1, together with p62 and LC3B, from the cytoplasm; when this export step is interrupted, KEAP1 is retained intracellularly and the NRF2 response to oxidative challenge is dampened. We emphasize that these experiments establish co-variation between export and NRF2 output rather than a causal chain, because dominant-negative RAB27A perturbs membrane traffic broadly; this constraint is addressed explicitly in the limitations below.

## Discussion

Using endogenous HiBiT tagging, we showed that p62 is constitutively secreted. The release of this protein is quantitatively comparable with that of LC3B, occurs predominantly in membrane-enclosed carriers recovered from the 10,000 × *g* pellet, and strongly increases when lysosomal degradation is blocked. Genetic dissection showed that the BafA1-inducible component requires ATG5 and FIP200, whereas the constitutive component does not. Furthermore, a non-sedimentable, detergent-independent pool indicates that some p62 leaves the cell without a surrounding membrane. p62 therefore exits cells via multiple routes.

Our data are compatible with four non-exclusive routes (Figure 5L). First, an autophagy-dependent route in which LC3-decorated membranes capture p62 and are diverted to the plasma membrane instead of the lysosome is supported by the membrane-enclosed nature of the secreted protein (Figures 2C and 3C). This is further supported by the loss of the BafA1-inducible component in ATG5 KO and FIP200 KO cells (Figures 4A and 4B), and the corresponding accumulation of LC3-II upon BafA1 treatment (Figure 1B). This route is also the most consistent with the LC3-dependent EV loading and secretion (LDELS) paradigm and with secretory autophagy during lysosome inhibition (Leidal and Debnath, 2020; Leidal et al., 2020; Solvik et al., 2022) and with the recent description of autophagy-derived EVs as a class distinct from canonical exosomes (Mao et al., 2025). LC3 can be conjugated to single-membrane endosomal and phagosomal compartments independently of the canonical pathway, as is the case in LC3-associated phagocytosis (Heckmann and Green, 2019). Therefore, an analogous non-canonical conjugation event could deliver p62 to a secretory compartment and would not be captured by our ATG5/FIP200 epistasis. The soluble N-ethylmaleimide-sensitive factor attachment protein receptor (SNARE) requirements also appear to differ from those of the interleukin (IL)-1β route. In particular, combined depletion of syntaxin 3 and syntaxin 4, which are plasma membrane Qa-SNAREs that complete IL-1β secretion downstream of tripartite motif containing 16 and vesicle-trafficking protein SEC22b (Kimura et al., 2017), did not reduce p62 release (K.S. and E.M., unpublished data). This experiment did not include a cargo that depends on these SNAREs as an internal positive control. Therefore, the study does not establish whether the fusion step that serves secretory autophagy of IL-1β is used by p62. This is discussed in the study limitations.

The second route is an autophagy-independent route that yields membrane-enclosed p62, most plausibly through intraluminal vesicle formation in the endosomal system (Figure 4A). The third is secretion of p62 as part of a higher-order receptor assembly. The selective reduction of BafA1-dependent secretion in the PB1 mutant K7A/D69A (Figure 5K) shows that oligomerization is required, whereas the absence of an effect of the L341V LIR variant shows that a direct p62–LC3 contact is not required. This mirrors the requirement for self-oligomerization but not LC3 binding in p62 targeting the autophagosome formation site (Itakura and Mizushima, 2011). This is most simply explained if other receptors in the same condensate (NBR1, TAX1BP1) supply the LC3 contact. The fourth is the release of non-enveloped p62, which may reflect lysosomal exocytosis or the escape of material after partial degradation (Figure 3C). In this study, these components are defined by their genetic and biochemical requirements rather than by the machinery that executes them. The fusion machinery that serves each route, which has been resolved for a subset of unconventional secretion events (Kimura et al., 2017), including UBB^+1^ secretion (Wagh and Glickman, 2025), lies outside the scope of the current study and is considered a limitation.

A domain analysis is informative in terms of what is excluded. Of the seven variants tested, only the oligomerization-deficient mutant exhibited altered secretion (Figure 5K). Neither the KIR phosphorylation status (S349A, S349E, or T350A), UBA phosphomimicry (S403E/S407E), nor UBA integrity (I431A) had a measurable effect. Therefore, the determinant of export is the assembly state of p62 rather than any single protein–protein interface. This is consistent with a model in which the unit of capture is a p62 condensate rather than an individual p62 molecule (Jakobi et al., 2020; Komatsu, 2022; Sun et al., 2018).

Extending the analysis to KEAP1 links this transport to redox signaling. In particular, KEAP1 was exported into the same membrane-enclosed carriers, and its release, similar to that of p62, increased upon lysosomal inhibition (Figure 6E). Disruption of p62 binding did not reduce KEAP1 secretion. This indicated that KEAP1 is not exported simply as a p62 cargo. The same mutations increased intracellular KEAP1 accumulation (Figure 6D), which suggests that binding-dependent mechanisms govern KEAP1 turnover, but not export. However, the mechanism by which KEAP1 reaches these carriers without binding to p62 was not resolved in this study. Client proteins are concentrated in p62 bodies through multivalent, low-affinity partitioning rather than through any single interface, and KEAP1 is among the clients that accumulate in them (Komatsu, 2022). Therefore, KEAP1 may still be enriched at the site of capture when the KIR–Kelch contact is lost. In addition, the Kelch pocket mutated in this study is the same surface that engages the ETGE motif of NRF2 and other Kelch-binding proteins. Therefore, an indirect route in which KEAP1 is delivered as part of a substrate-bound KEAP1–CUL3 complex is equally compatible with our data. Notably, blocking RAB27-dependent release reduced the secretion of LC3B, p62, and KEAP1, and lowered NRF2 reporter activity and induction of the endogenous NRF2 target HO-1 under oxidative challenge (Figures 7A–7D). Continuous vesicular export may therefore constitute a route of KEAP1 disposal that operates in parallel with p62-mediated sequestration. A parsimonious interpretation of the RAB27A data is that interrupting this route increases the pool of intracellular KEAP1 available to target NRF2 for degradation, although our experiments do not exclude contributions from other RAB27A-dependent trafficking events, and we make no claim that KEAP1 export is the dominant determinant of NRF2 output. With that reservation, the data are compatible with a model in which the secretory capacity of the cell, and not only its degradative capacity, feeds into the oxidative stress response.

The HiBiT KI strategy resolves several long-standing practical obstacles in this area. It measures the endogenous protein, requires no antibody, spans the concentration range between the intracellular and extracellular pools in a single assay, and reports membrane topology directly through detergent dependence. Applied to p62, it established that a core autophagy receptor is exported by multiple routes, that its export tracks the status of the degradative system, and that its binding partner KEAP1 is exported with it.

We consider the generality of this design to be the most transferable outcome of the present work. The quantitative study of unconventional secretion has been constrained less by biological hypotheses than by measurement: immunoblotting of conditioned medium lacks the sensitivity to detect low-abundance cytosolic cargo, vesicle purification recovers material with variable and rarely calibrated efficiency, and the overexpression normally used to compensate alters the very assembly behavior that determines sorting. Endogenous HiBiT tagging addresses these three problems simultaneously. Because the tag is 11 residues and is inserted at the native locus, protein abundance, turnover, and condensate behavior are preserved; because the readout is enzymatic complementation, the same standard curve covers the intracellular pool and the far smaller extracellular pool without fractionation-dependent normalization; and because LgBiT is membrane-impermeant, a paired measurement with and without detergent converts an otherwise separate topology experiment into a two-well assay. The consequence is that a secretion rate can be expressed as a fraction of the total cellular pool for each protein, which makes different cargoes directly comparable within one experiment, as illustrated here for p62, LC3B, and TOMM20.

The same design should apply to any low-abundance cytosolic protein whose extracellular fate is suspected but has been difficult to quantify, including other selective autophagy receptors (NBR1, TAX1BP1, optineurin, NDP52), the leaderless cytokines and RNA-binding proteins reported as secretory autophagy or LDELS cargo, and aggregation-prone proteins whose intercellular transfer is of interest in neurodegeneration. Because the assay is plate-based, it is also compatible with the combinatorial or acute loss-of-function screens that, as discussed above, will be required to assign fusion machinery to each route. We therefore regard the panel of matched knock-in lines described here as a resource as much as a result.

### Limitations of this study

The study has some limitations. First, we did not identify the membrane fusion machinery responsible for p62 release. Assigning specific SNAREs to the routes summarized in Figure 5L would convert components that are currently distinguished operationally into molecularly defined pathways, and would provide the selective loss-of-function tools that a study of parallel secretory routes requires. We depleted candidate SNAREs implicated in secretory autophagy and EV release using small-interfering RNA but did not obtain a condition that reproducibly reduced p62 secretion. This included a combined knockdown of syntaxin 3 and syntaxin 4, which are the plasma membrane Qa-SNAREs required for IL-1β secretion by secretory autophagy. This knockdown did not alter p62 release (K.S. and E.M., unpublished data). These experiments did not include a cargo that depends on these SNAREs as an internal positive control. Therefore, we could not formally distinguish between incomplete depletion and the genuine independence of this fusion step. We regard this as a technical rather than a conceptual outcome. SNAREs of the secretory and endolysosomal compartments are functionally redundant. Specifically, the residual protein remaining after partial knockdown is frequently sufficient to support fusion. Given that p62 leaves the cell by more than one route, blocking a single fusion step may redirect cargo into a parallel route rather than retain it intracellularly. The identification of SNAREs that serve secretory autophagy required a dedicated effort in previous studies (Kimura et al., 2017), and a comparable effort (combinatorial depletion or acute inactivation that outpaces compensation) will be needed here. Therefore, we have described the routes based on the requirements we could test and make no claim about the fusion machinery that serves them. However, once the machinery is known, the contribution of each route will be clarified.

Second, the RAB27A experiments established that vesicular export and NRF2 output co-vary. However, these experiments did not establish that KEAP1 export is the dominant determinant of NRF2 activity. Dominant-negative RAB27A broadly perturbs membrane traffic, and we therefore restrict our claim to what the measurements support: interrupting a RAB27A-dependent export step retains KEAP1, p62, and LC3B intracellularly and lowers both NRF2 reporter output and induction of the endogenous target HO-1 under oxidative challenge. Whether extracellular KEAP1-containing carriers themselves are signaling-competent in recipient cells requires further investigation.

Finally, the secreted material was assigned to operationally defined fractions. In line with MISEV2023 (Welsh et al., 2024), we named the fractions according to their isolation procedure rather than by EV subtype, and the conclusions relied on the sedimentation behavior and detergent-dependent accessibility of the HiBiT tag rather than on subtype assignment. Secretion rates are also expressed as a fraction of the total HiBiT signal for each protein because the luminescence yield of the reconstituted HiBiT–LgBiT complex may differ between fusion proteins, and comparisons of secretion rates between p62, LC3B, and TOMM20 are relative rather than absolute.

## Materials and methods

### Key resources table

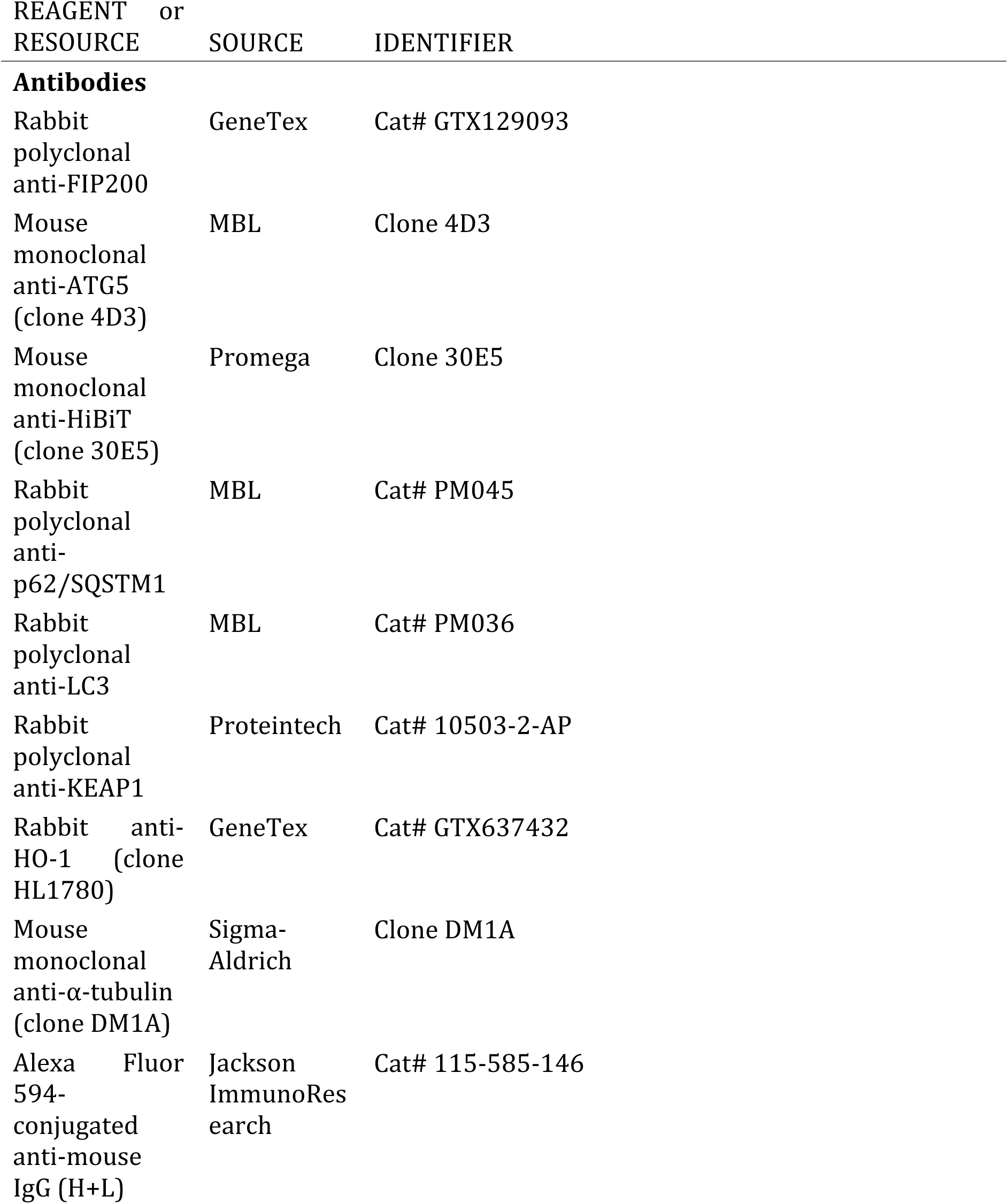
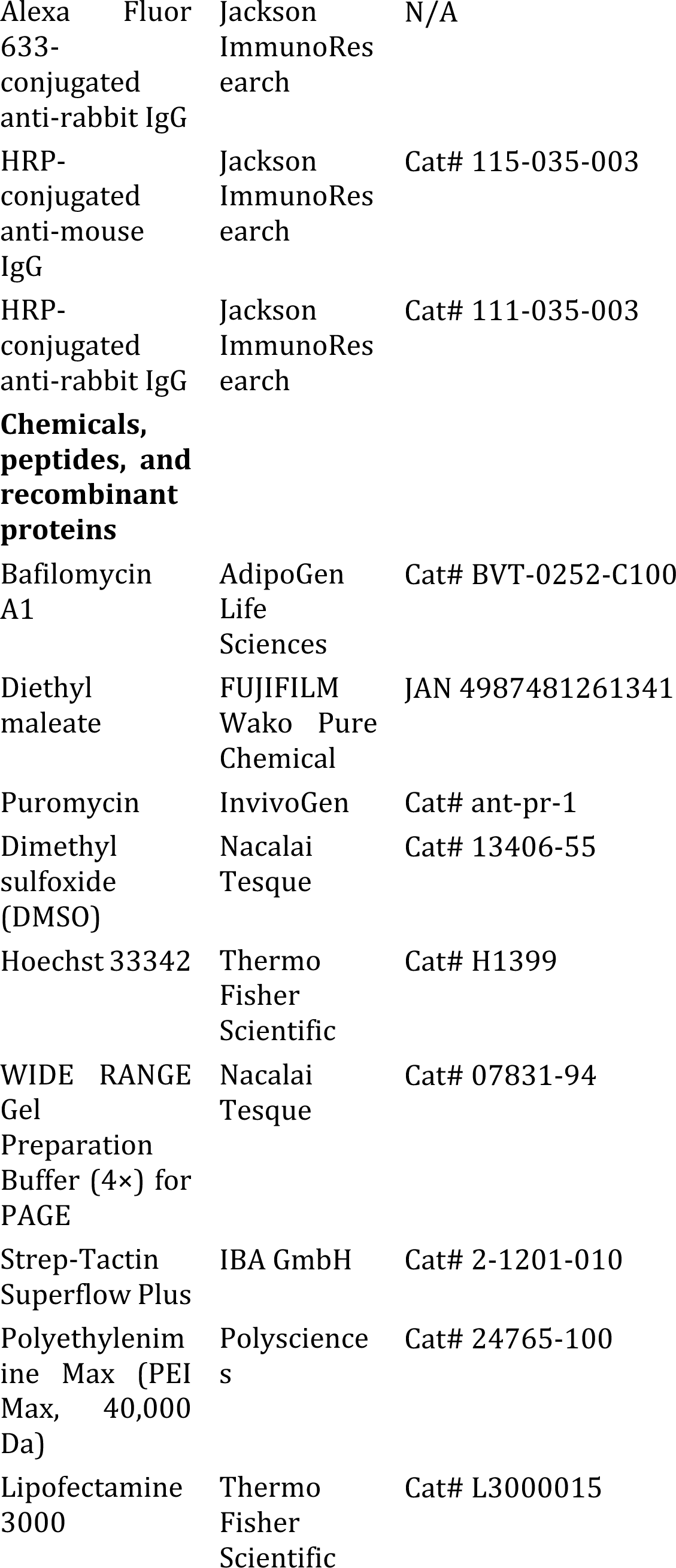
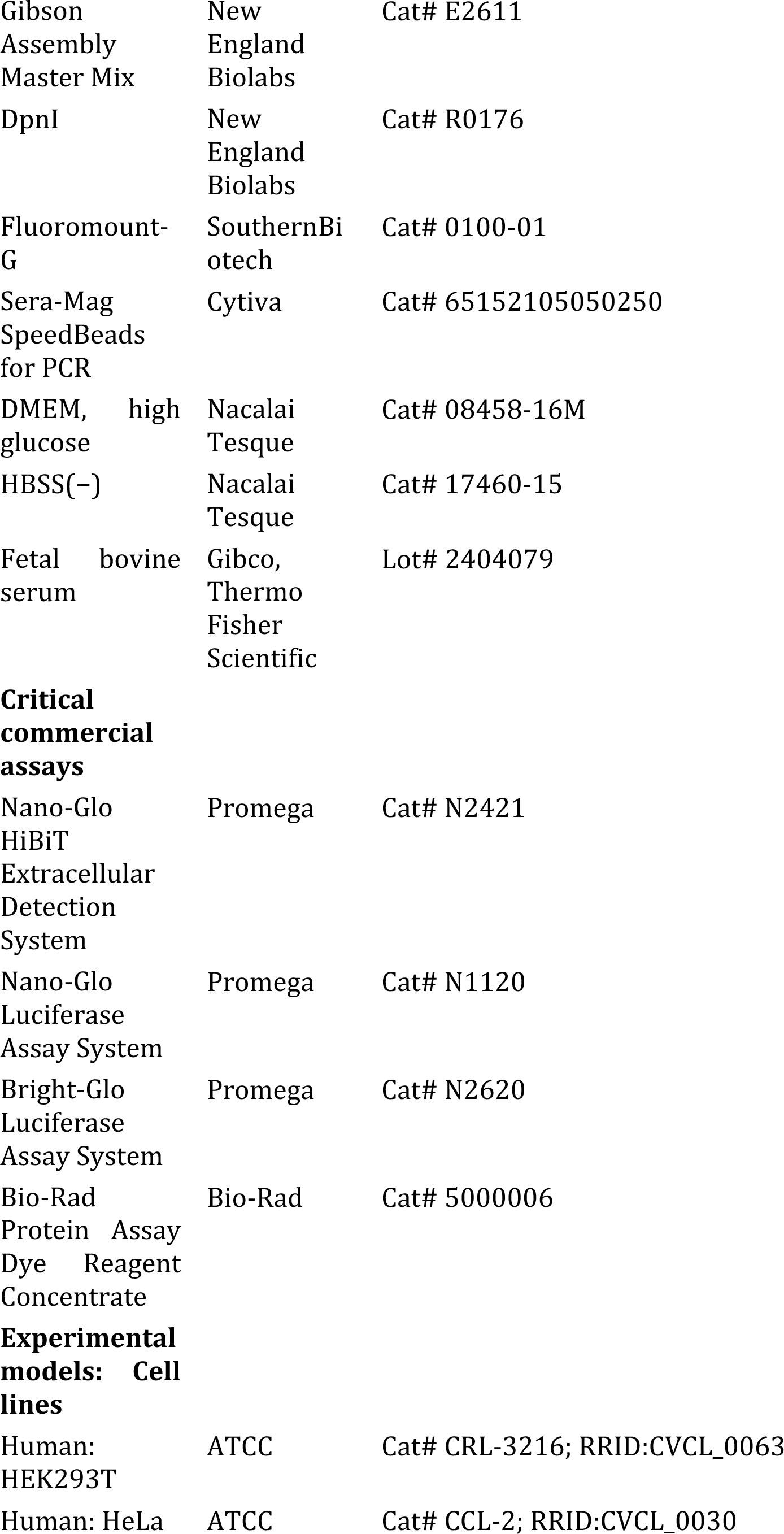
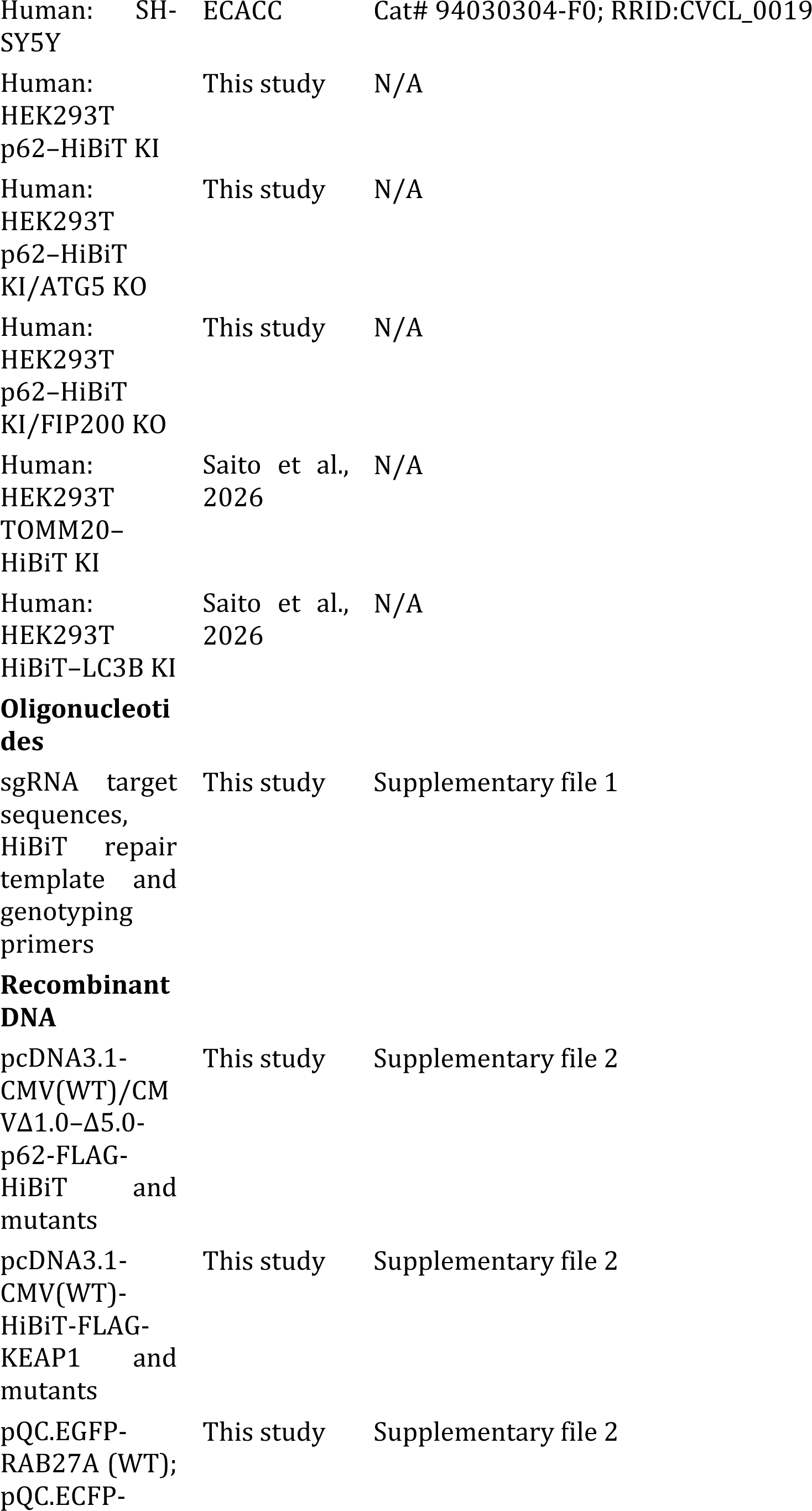
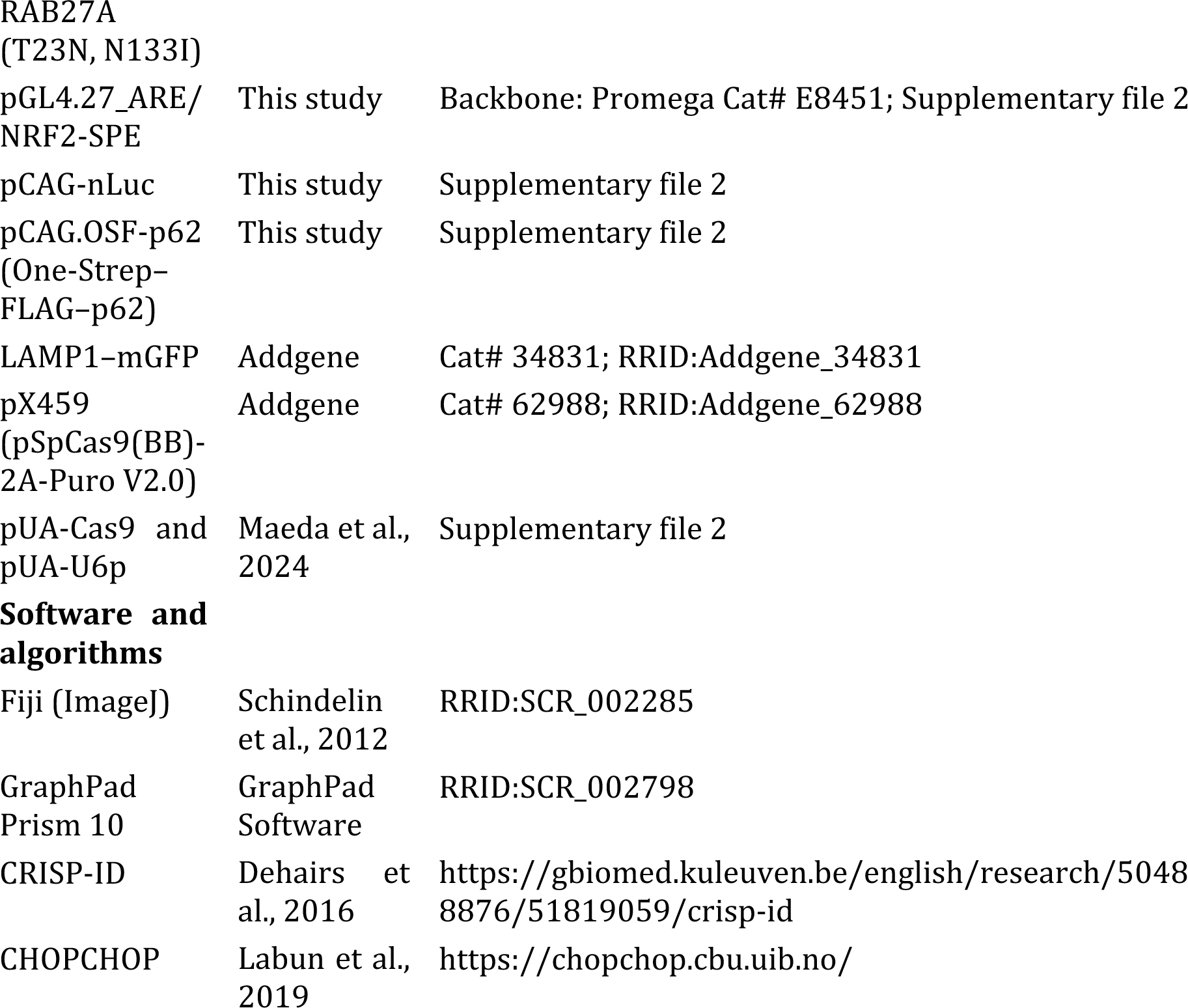

### Cell lines

HEK293T (human embryonic kidney, female; ATCC Cat# CRL-3216), HeLa (human cervical adenocarcinoma, female; ATCC Cat# CCL-2) and SH-SY5Y (human neuroblastoma, female; ECACC Cat# 94030304-F0) cells were maintained in Dulbecco’s modified Eagle’s medium, high glucose (DMEM; Nacalai Tesque) supplemented with 10% fetal bovine serum (lot 2404079; Gibco, Thermo Fisher Scientific), 100 U/mL penicillin (Nacalai Tesque) and 100 μg/mL streptomycin (Nacalai Tesque) at 37°C under 5% CO2. Genome-edited lines generated in this study were cultured under identical conditions. The cell lines were not authenticated further and were routinely confirmed to be free of mycoplasma contamination. No human or animal participants were included in this study.

### Plasmid construction and transfection

All plasmids were constructed by restriction ligation or Gibson assembly and are listed in Supplementary file 2. For restriction ligation, PCR-amplified inserts were digested with the appropriate restriction enzymes and ligated into multiple cloning sites of the corresponding vector. For Gibson assembly, vector and insert were amplified with primers carrying ≥15-nucleotide homology arms and joined using Gibson Assembly Master Mix (New England Biolabs) together with DpnI (New England Biolabs); reactions were incubated at 37°C for 30 min followed by 50°C for 60 min. The plasmids were introduced into HEK293T, HeLa, and SH-SY5Y cells using polyethylenimine (PEI Max, 40,000 Da; Polysciences) or Lipofectamine 3000 (Thermo Fisher Scientific) according to the manufacturer’s instructions.

### CRISPR-Cas9 genome editing

p62–HiBiT knock-in HEK293T cells were generated by CRISPR-Cas9-mediated insertion of the HiBiT coding sequence into the frame at the C-terminus of endogenous p62/SQSTM1. Using these cells as parental lines, *ATG5* or *RB1CC1* (FIP200) was disrupted to generate p62–HiBiT KI/ATG5 KO and p62–HiBiT KI/FIP200 KO cells. Guide RNA target sequences were designed using CHOPCHOP (Labun et al., 2019) and are listed, together with the repair template and genotyping primers, in Supplementary file 1.

Editing was performed with pX459 (pSpCas9(BB)-2A-Puro V2.0; Addgene, Cat# 62988 (Ran et al., 2013)) or with the pUA-Cas9 and pUA-U6p vector pairs (Maeda et al., 2024), which express Cas9 together with sgRNA driven by a U6 promoter and a puromycin resistance cassette. For knock-in, a repair template containing the HiBiT sequence flanked by 33-nucleotide homology arms was cotransfected. Transfected cells were selected with puromycin (1.25 μg/mL; InvivoGen) and single clones were isolated by limiting dilution. Each clone was validated using PCR amplification and sequencing of the target region.

### Validation of genome editing

The edited clones were verified by Sanger sequencing and immunoblotting. Genomic DNA was prepared by suspending the cell pellets in ddH2O, and the region flanking the target site was amplified by PCR using the primers listed in Supplementary file 1. The products were verified using agarose gel electrophoresis, purified using magnetic beads (Sera-Mag for PCR; Cytiva), and used as templates for Sanger sequencing. Sequence traces were deconvoluted using CRISP-ID (Dehairs et al., 2016) to confirm HiBiT knock-in and *ATG5* or *RB1CC1* disruption. For protein level validation, cells were lysed in SDS sample buffer (125 mM Tris-HCl [pH 6.8], 4% SDS, 20% glycerol, 0.01% bromophenol blue, and 10% 2-mercaptoethanol), resolved by SDS-PAGE, and immunoblotted. p62–HiBiT was detected with both anti-HiBiT and anti-p62 antibodies, and the loss of ATG5 and FIP200 was confirmed with the respective antibodies.

### Autophagic flux assay

Autophagic flux was assessed by immunoblotting of the conversion of LC3-I to LC3-II. Starvation was induced by replacing the medium with Hanks’ balanced salt solution (HBSS(−); Nacalai Tesque) for 6 h. Lysosomal function was inhibited with 100 nM bafilomycin A1 (BafA1; AdipoGen Life Sciences) dissolved in DMSO (Nacalai Tesque); DMSO alone served as vehicle control. Flux was evaluated as the increase in LC3-II levels under starvation and BafA1 treatment.

### Isolation of extracellular fractions

Extracellular fractions were isolated by differential centrifugation, as previously described (Saito et al., 2026). Conditioned medium was cleared of cells and debris by centrifugation at 500 × *g* for 5 min and 1,200 × *g* for 5 min at 4°C. The cleared supernatant was centrifuged at 10,000 × *g* for 30 min to recover a large EV-enriched pellet (10k ppt). The resulting supernatant (10k sup) was centrifuged at 100,000 × *g* for 70 min to recover the small EV-enriched pellet (100k ppt), and the final supernatant was collected as the non-sedimentable fraction (100k sup). Fraction nomenclature and reporting followed the MISEV2023 recommendations (Welsh et al., 2024). Operationally defined fractions were referred to by their centrifugal parameters rather than by biogenesis-implying terms.

### HiBiT complementation assay

HiBiT activity was measured using a Nano-Glo HiBiT Extracellular Detection System (Promega) (Dixon et al., 2016). The LgBiT solution was prepared by mixing the HiBiT extracellular buffer, LgBiT protein, and substrate at a ratio of 100:1:2. An equal volume of LgBiT solution was added to each sample (10 μL) in a 384-well white plate (Greiner Bio-One), and luminescence was recorded with a Varioskan LUX plate reader (Thermo Fisher Scientific) using a 1 s integration time.

To assess membrane enclosure, parallel measurements were performed with LgBiT solution containing 0.1% Triton X-100 (detergent +) and without detergent (detergent −); signal detected only in the presence of detergent was taken to indicate protein enclosed within a sealed membrane.

### NRF2 reporter assay

SH-SY5Y cells were seeded in 12-well plates at a density of 3 × 10^5^ cells/well and transfected with PEI. Each well received 1,000 ng of pQC.EGFP-RAB27A (wild-type) or pQC.ECFP-RAB27A (T23N or N133I), 990 ng of pGL4.27_ARE/NRF2-SPE, and 10 ng of pCAG-nLuc as an internal control. Cells were treated with or without 500 μM diethyl maleate (DEM) for 6 h and harvested. Firefly luciferase activity was measured using a Bright-Glo Luciferase Assay System (Promega) and normalized to NanoLuc activity, which was measured using a Nano-Glo Luciferase Assay System (Promega).

### Immunoblotting and antibodies

Proteins were separated on polyacrylamide gels cast with WIDE RANGE Gel Preparation Buffer (4×) for PAGE (Nacalai Tesque, Kyoto, Japan), and transferred to PVDF membranes. The primary antibodies used were as follows: anti-FIP200, 1:1,000; anti-ATG5, 1:1,000; anti-HiBiT, 1:10,000 for immunoblotting and 1:1,000 for immunofluorescence; anti-LC3, 1:1,000; anti-p62, 1:1,000; anti-KEAP1, 1:10,000; anti-HO-1, 1:500; and anti-α-tubulin, 1:1,000. Horseradish peroxidase-conjugated anti-mouse IgG and anti-rabbit IgG (both 1:3,000; Jackson ImmunoResearch) were used as secondary antibodies.

### Protein quantification

The protein concentration was determined using a Bio-Rad Protein Assay (Bio-Rad). The reagent was diluted fivefold with Milli-Q water. Cells were lysed in lysis buffer (1% Triton X-100, 50 mM Tris-HCl [pH 7.4], 150 mM NaCl, protease inhibitors), and 1 μL of sample was mixed with 100 μL of reagent in a 96-well plate. The absorbance was measured at 595 nm (Varioskan LUX). The concentrations were calculated using a standard curve prepared using bovine serum albumin.

### Immunofluorescence microscopy

Cells were grown on coverslips, transfected as indicated, and fixed in 4% paraformaldehyde for 10 min at room temperature. After washing with phosphate-buffered saline (PBS), the cells were permeabilized with 0.1% Triton X-100 in PBS for 10 min. Blocking and primary antibody incubation were performed in PBS containing 10% FBS for 1 h at room temperature, together with Hoechst 33342 (1:1,000; Thermo Fisher Scientific) for nuclear staining. After washing, the cells were incubated with Alexa Fluor 594-conjugated anti-mouse IgG or Alexa Fluor 633-conjugated anti-rabbit IgG (both 1:1,000; Jackson ImmunoResearch) for 1 h at room temperature, washed, and mounted in Fluoromount-G (Southern Biotech). Images were acquired using an FV3000 confocal laser-scanning microscope (EVIDENT).

### Affinity purification

Proteins expressed in the HEK293T cells were purified using Strep-Tactin resin. Cells were lysed in lysis buffer, incubated on ice, and cleared by centrifugation at 4°C for 10 min. The supernatant was rotated with Strep-Tactin Superflow Plus (IBA GmbH) for 2 h at 4°C. The resin was recovered and washed several times with wash buffer (0.1% Triton X-100, 150 mM NaCl, and 20 mM Tris-HCl [pH 7.4]). The bound proteins were eluted by adding 2× sample buffer and heating at 95°C for 5 min. Total cell lysate controls were prepared by mixing an aliquot of the same lysate with an equal volume of 2× sample buffer and heating at 95°C for 5 min.

### Quantification and statistical analysis

Immunofluorescence images and immunoblot band intensities were quantified using the Fiji software (ImageJ) (Schindelin et al., 2012). HiBiT activity in the cell fractions was normalized to that of the total protein. The secretion rate (%) was calculated as the HiBiT activity in the indicated extracellular fraction divided by the sum of the activities in that fraction and the corresponding cell fraction. For graphical presentation, the secretion rates were normalized to the indicated reference conditions in each experiment, which were set to 1. NRF2 activity was calculated as firefly luciferase luminescence divided by NanoLuc luminescence.

Statistical analyses were performed using the GraphPad Prism 10 software (GraphPad Software). The two groups were compared using an unpaired two-tailed Student’s t-test. Multiple groups were compared against the control group using one-way ANOVA with Dunnett’s post hoc test. Two-factor designs were analyzed using two-way analysis of variance (ANOVA) followed by Tukey’s multiple comparison test. Where differences between groups were expected to be multiplicative, as for HiBiT luminescence in Figures 1C and 2B, values were log-transformed before analysis, and background values from parental cells were excluded from the ANOVAs. In Figure 1C, the effects of BafA1 and HBSS were tested within each genotype using Šídák’s multiple comparison test, and p62–HiBiT KI and parental WT cells under control conditions were compared by lognormal Welch’s t test. Data are presented as mean ± SD of n = 3 technical replicates; each experiment was performed independently at least twice with comparable results, and the figures show a representative experiment. The statistical test used, value of n, and definition of the error bars are stated in each figure legend. Significance is indicated by *p < 0.05, **p < 0.01, ***p < 0.001, and ****p < 0.0001. ns, not significant.

## Data availability

All data generated or analysed during this study are included in the manuscript and supporting files. Source data for all quantified figure panels are available from the corresponding author on request. This study does not report original code.

## Materials availability

Plasmids and cell lines generated in this study are available from the corresponding author on request without restriction, subject to a completed material transfer agreement.

## Acknowledgements

This work was supported by JSPS KAKENHI (grant numbers 22H02873, 22H00553, 26K02144, and 26K18636), the Takeda High-Risk Emerging Infectious Diseases Research Grant, the Daiichi Sankyo Foundation of Life Science, the NOVARTIS Foundation (Japan) for the Promotion of Science, The Naito Foundation, the MEXT J-PEAKS (Program for Forming Japan’s Peak Research Universities) Grant Number [JPJS00420240013], and Hirosaki University Grants for Next-Generation Key Research Areas.

## Author contributions

Conceptualization: E. M. Investigation: K. S. and T. T. Methodology: K. S. Validation: K. S., T. T., and M. A. Formal analysis: K. S. and T. T. Visualization: K. S. and M. A. Resources: K. S. Writing – Original Draft: K. S. and E. M. Writing – Review & Editing: M. A. and E. M. Supervision: M. A. and E. M. Funding Acquisition: M. A. and E. M.

## Use of generative AI

During the preparation of this work, the authors used Claude (Anthropic) in order to improve the readability and language of the manuscript. After using these tools, the authors reviewed and edited the content as needed and take full responsibility for the content of the publication.

## Supplementary files

**Supplementary file 1.**
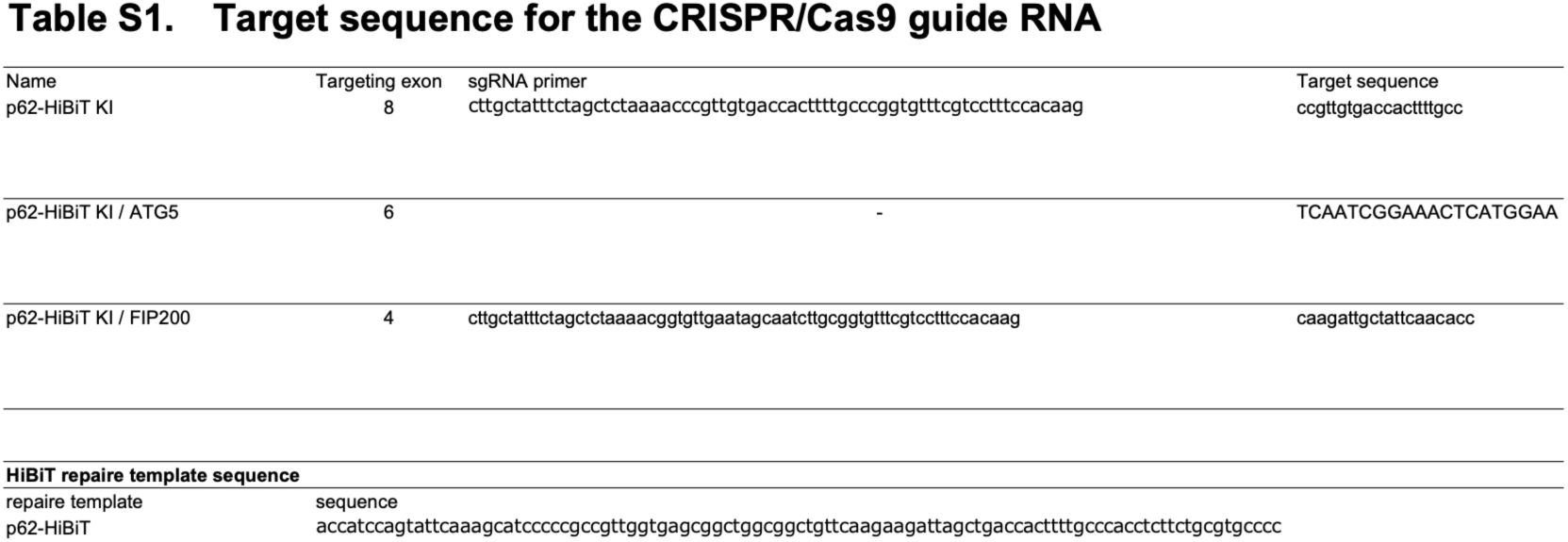
Oligonucleotides used in this study.

**Supplementary file 2.**
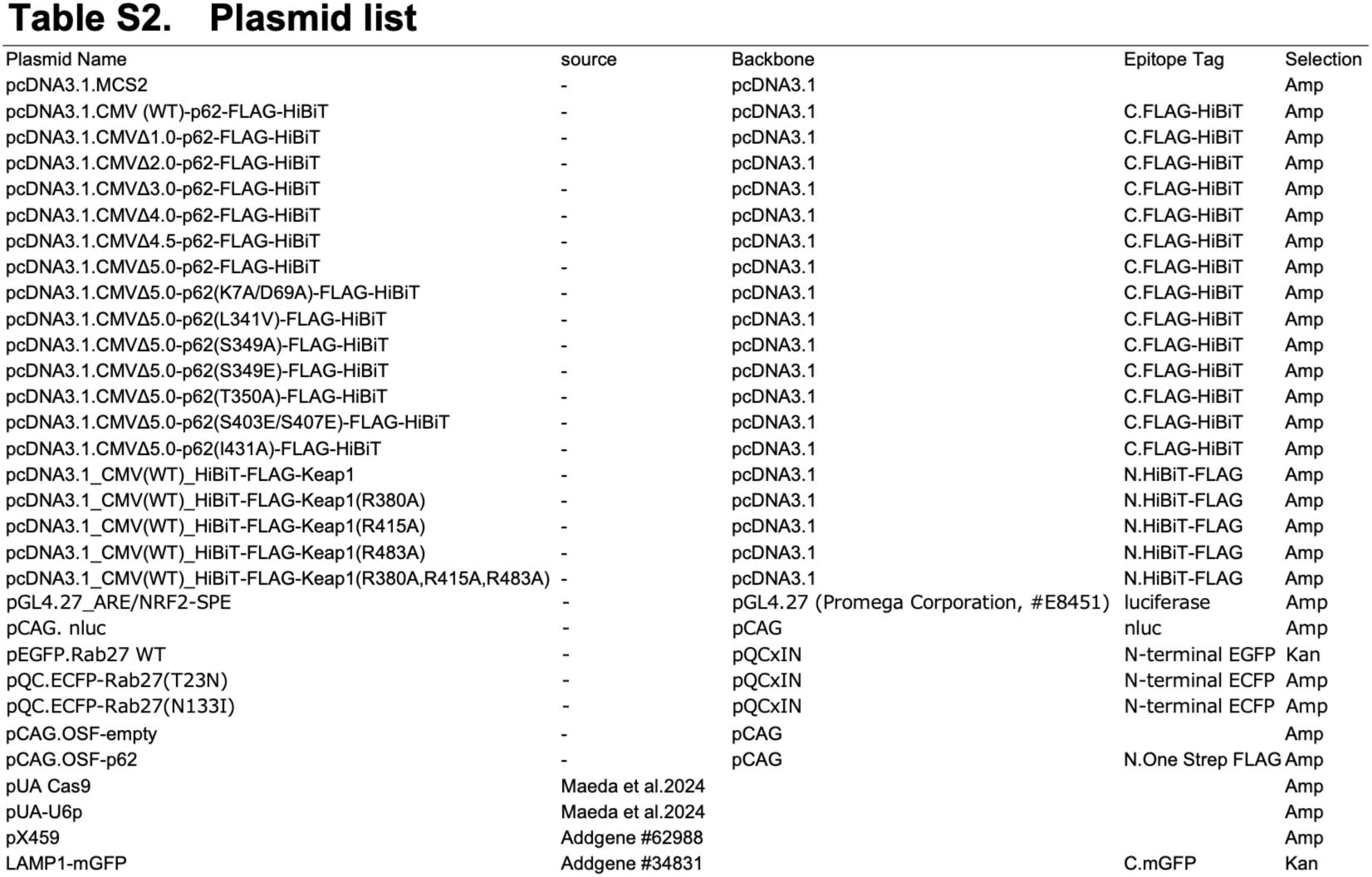
Plasmids used in this study.

